# Oscillatory and aperiodic neural activity jointly predict language learning

**DOI:** 10.1101/2020.03.10.984971

**Authors:** Zachariah R. Cross, Andrew W. Corcoran, Matthias Schlesewsky, Mark. J. Kohler, Ina Bornkessel-Schlesewsky

## Abstract

Memory formation involves the synchronous firing of neurons in task-relevant networks, with recent models postulating that a decrease in low frequency oscillatory activity underlies successful memory encoding and retrieval. However, to date, this relationship has been investigated primarily with face and image stimuli; considerably less is known about the oscillatory correlates of complex rule learning, as in language. Further, recent work has shown that non-oscillatory (1/*f*) activity is functionally relevant to cognition, yet its interaction with oscillatory activity during complex rule learning remains unknown. Using spectral decomposition and power-law exponent estimation of human EEG data (17 females, 18 males), we show for the first time that 1/*f* and oscillatory activity jointly influence the learning of word order rules of a miniature artificial language system. Flexible word order rules were associated with a steeper 1/*f* slope, while fixed word order rules were associated with a shallower slope. We also show that increased theta and alpha power predicts fixed relative to flexible word order rule learning and behavioural performance. Together, these results suggest that 1/*f* activity plays an important role in higher-order cognition, including language processing, and that grammar learning is modulated by different word order permutations, which manifest in distinct oscillatory profiles.

## Introduction

Memory supports many essential cognitive functions, from learning the distinction between semantic categories (e.g., animal vs. human) to complex (motor) sequences, such as learning how to drive a car or speak a new language. However, while a broad literature has related neural oscillatory dynamics (i.e., [de]synchronisation of neural populations) to the encoding and retrieval of images and words (e.g., Parish, Hanslmayr, & Bowman, 2018), considerably less is known about oscillatory activity during the encoding and retrieval of complex sequences, such as in language (cf. de Diego-Balaguer, Fuentemilla, & Rodriguez-Fornells, 2011; Kepinska, Pereda, Caspers, & Schiller, 2017). Further, the few studies examining sequence and artificial language learning report mixed findings relative to (episodic) word and image paradigms: while alpha/beta desynchronisation in the human EEG predicts encoding of words and images (e.g., Griffiths, et al., 2019), alpha/beta and theta synchronisation is associated with sequence (Crivelli-Decker et al., 2018) and language learning (e.g., Kepinska et al., 2017). This apparent inconsistency might be accounted for by stimulus heterogeneity; however, another possible source of divergence may lie in the mixture of oscillatory power with aperiodic activity (Ouyang et al., 2020; Wen & Liu, 2016), which has not been addressed in studies on the neural basis of complex rule learning to date.

Electrophysiological brain activity exhibits a 1/*f*-like power distribution, which is often removed from the signal to isolate transient task-related oscillations (Donoghue et al., 2020; He, 2014; Lendner et al., 2020). However, this aperiodic component has recently been implicated in a variety of higher-order cognitive computations (Fellner et al., 2019), partially explaining individual differences in theta activity during memory encoding and recall performance (Sheehan et al., 2018), and processing speed over and above that of alpha activity (Ouyang et al., 2020). Work on prediction during language has also shown that the aperiodic slope – but not oscillatory activity – influences the N400 event-related potential and performance accuracy (Dave et al., 2018).

These findings suggest that aperiodic brain activity plays a critical functional role in the neurobiology of cognition (He et al., 2010); however, it is currently unknown whether oscillatory and aperiodic activity interact during memory encoding of information beyond single words and images, such as rule-based sequence learning, and whether any such interaction influences behavioural outcomes. Clarifying the (separable) roles of oscillatory and aperiodic components of the EEG power spectrum may also bridge diverging results reported in studies using image and word stimuli and artificial grammar paradigms, lending support to the idea that neural oscillations differentially contribute to memory formation.

To better characterise the neural mechanisms underlying complex rule learning, we examined fluctuations in delta, theta, alpha and beta power during an artificial language learning task. We also modelled the interaction between oscillatory and aperiodic activity to characterise how patterns of (de)synchronisation and aperiodic fluctuation influence the generalisation of different word order rules characteristic of many natural languages. Healthy young adults learned the artificial miniature language Mini Pinyin (Cross, Zou-Williams, Wilkinson, Schlesewsky, & Bornkessel-Schlesewsky, 2020a) without explicit instruction and then completed a sentence judgement task. Critically, participants – who were native monolingual English speakers – learned fixed and flexible word order rules: fixed word order sentences contained temporal- or sequence-based rules, while flexible word order sentences involved non-adjacent dependencies, likely relying on more associative-than sequence-based memory processing mechanisms (Cross et al., 2018). From this perspective, we were able to probe different learning and memory mechanisms that are involved in sentence comprehension (Bornkessel-Schlesewsky, Schlesewsky et al., 2015). For example, native English speakers typically rely on word-order-based cues for sentence comprehension, whereas speakers of Mandarin Chinese or the Australian language Jiwarli rely more strongly on cues other than word order, such as case marking and/or semantic information, including animacy (Austin, 2001; Bates et al., 2001; Bornkessel-Schlesewsky et al., 2011).

We recorded EEG during the learning task, implementing generalised additive and linear mixed-effects regression analyses to model dynamic changes in oscillatory and aperiodic activity during the learning of the fixed and flexible word order rules. We also modelled learning-related oscillatory and aperiodic activity to predict subsequent behavioural performance on the sentence judgement task, quantified as the sensitivity index d’.

## Method

### Participants

Data from 36 right-handed healthy, monolingual, native English-speakers were used from a study examining the effect of sleep on language learning (Cross et al., 2021). This sample size was based on previous EEG research examining the neural correlates of higher-order language learning (de Diego-Balaguer et al., 2011; Kepinska et al., 2017; Mueller et al., 2005). One participant was excluded from analysis for not having electroencephalography recorded during the sentence learning task due to experimenter error. The final sample size was 35 (M_age_ = 25.3, *SD* = 7.13; 17 female). Ethics approval was granted by the University of South Australia’s Human Research Ethics committee (I.D.: 0000032556).

### Stimuli and experimental design

Stimuli were based on the modified miniature language Mini Pinyin (for a detailed description of the language, see Cross et al., 2020a; see also Cross et al., 2021), which contains grammatical rules present in a number of natural languages (see Figure 1A and 1B for example sentence constructions and vocabulary items). Briefly, each sentence in Mini Pinyin contains two noun phrases and a verb, and each noun is associated with a different classifier: human nouns are preceded by *ge*, while animals, and small and large objects are preceded by *zhi, xi* and *da*, respectively.

**Figure 1.**
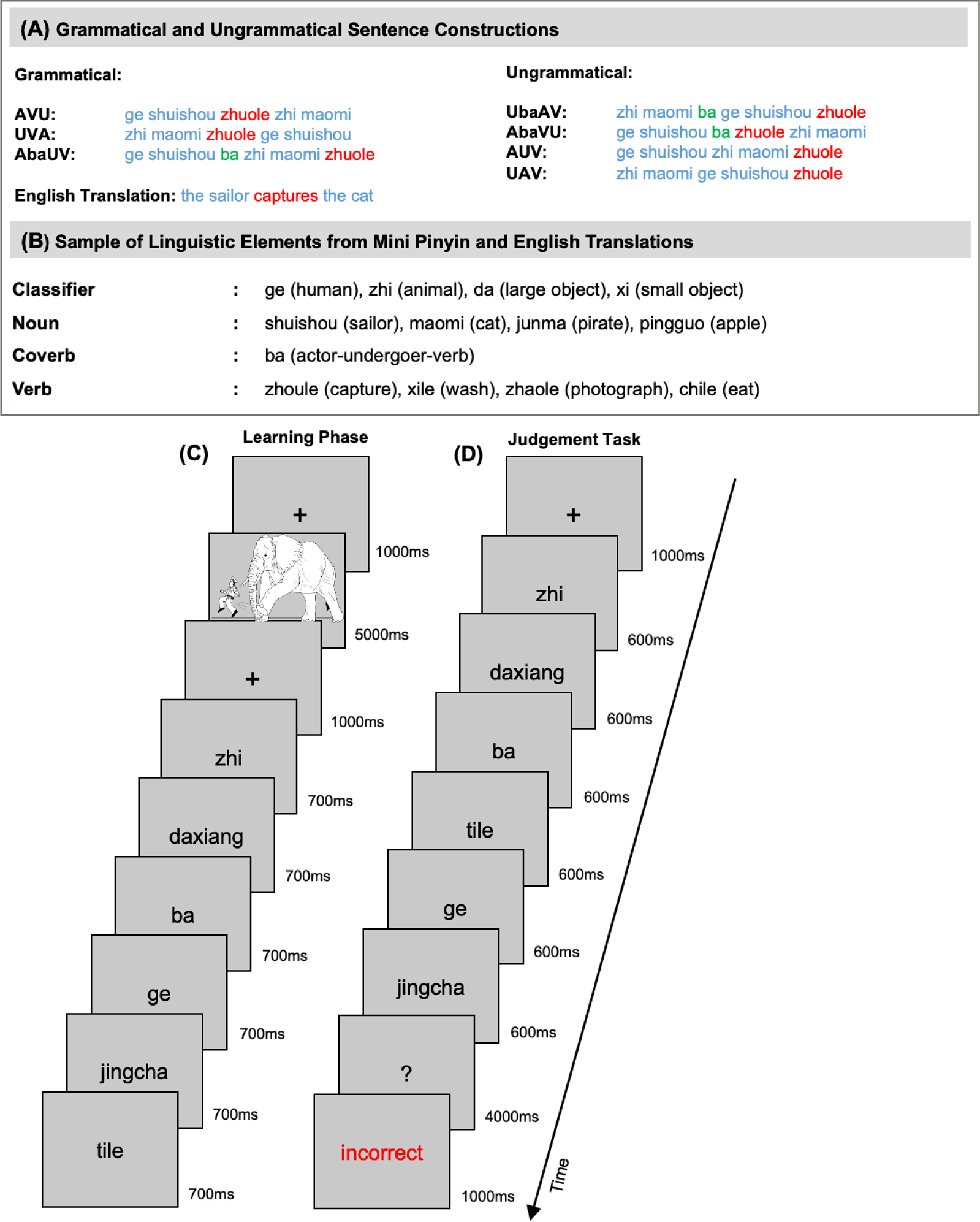
(A) Summary of the grammatical (left) and ungrammatical (right) sentence constructions. (B) A portion of linguistic elements used in the sentence examples provided in (A). (C) Schematic of the sequence of events occurring in the sentence learning phase. (D) Schematic of the sequence of events occurring in the sentence judgement task.

Mini Pinyin includes two main sentence types based on whether the sentence contains the coverbs *ba* and *bei*. Here we focus on the coverb *ba*: when *ba* is present, the sentence contains a fixed word order, in that the first noun phrase is invariably the Actor (the active, controlling participant in the action described by the sentence) and the sentence must be verb final. In this context, accurate sentence processing is dependent on the linear position of the words. When *ba* is not present, the sentence contains a flexible word order, in that the first noun phrase can either be the Actor or the Undergoer (the affected participant); however, the sentence must be verb-medial (i.e., verb must be positioned between the noun phrases). As such, accurate sentence interpretation is based more heavily on the animacy status of the noun phrases rather than word order. These manipulations are illustrated below, with a flexible word order shown in 1a and 1b and fixed (i.e., with the coverb *ba*) shown in 2:

(1) (a) ge yisheng dale da piqiu. (human) doctor hit (object) ball. “the doctor hits the ball.”
(b) da xianjiao chile zhi laoshu. (object) banana eat (animal) rat. “the rat eats the banana.”
(2) ge xiaofang ba da shubao liangle. (human) firefighter ba (object) bag measure. “the firefighter measures the bag.”

The experiment contained three phases: a vocabulary test, a learning phase, and a grammaticality judgement task. Approximately seven days before the sentence learning phase, participants received a paired picture-word vocabulary booklet containing the 25 nouns. Participants were required to learn the 25 nouns to ensure that they had a basic vocabulary. Prior to the learning and judgement tasks, participants completed the vocabulary test on a desktop computer by typing in translations of the nouns from Mini Pinyin to English. Only participants who attained a score > 90% were eligible to undertake the learning and judgement task phases of the experiment. All 35 participants achieved 90% accuracy on the vocabulary task and thus completed the main experimental session. Overall, 576 unique sentences (288 grammatical, 288 ungrammatical) were created and divided into two equivalent sets.

We focus here on a subset of sentence conditions to investigate the mechanisms underlying the learning of different word order permutations (for EEG analyses during the sentence judgement task, see Cross et al., 2021). While no explicit instructions were given to participants in regard to the structure of the miniature language, a picture was shown prior to each sentence illustrating an event occurring between two entities, which was then described in the subsequently presented sentence. The learning task contained four blocks with 128 grammatical picture-sentence pairs overall (96 of which were included in the subset analysed here) that were presented via rapid visual serial presentation. The subset contained a further 156 novel sentences (50% grammatical, 50% ungrammatical) that were presented during the judgement task, which occurred immediately after the learning phase. The remaining sentences were considered fillers. The ungrammatical sentences induced a violation at either the position of the Actor or verb in fixed word order sentences (e.g., Actor-ba-Verb-Undergoer [AbaVU] instead of AbaUV) or the position of the verb in flexible word order sentences (e.g., AUV instead of AVU; see Figure 1A for an illustration and full list of ungrammatical constructions).

During the learning phase, each picture was presented for 5000ms, while each corresponding sentence was presented on a word-by-word basis, with each word presented for 700ms with an inter-stimulus interval (ISI) of 200ms. Across the four blocks, each grammatical construction was presented 32 times, with stimuli pseudo-randomised such that no sentences of the same construction followed each other. During the judgement task, novel grammatical and ungrammatical sentences were presented word-by-word with a presentation time of 600ms and an ISI of 200ms. Participants responded via a button press to indicate whether the sentence conformed to the rules of Mini Pinyin. The assignment of grammatical/ungrammatical response buttons was counterbalanced across participants. Response time windows were presented for a maximum of 4000ms. Participants received feedback on whether their response was correct or incorrect (see Figure 1C and 1D for a schematic of the learning and judgement tasks, respectively). Both the learning and judgement tasks were created in OpenSesame (Mathot et al., 2012) and performed on a desktop computer.

### EEG recording and pre-processing

Participants’ EEG was recorded using a 32-channel BrainCap with sintered Ag/AgCl electrodes (Brain Products, GmbH, Gilching, Germany) mounted according to the extended International 10-20 system. The online reference was located at FCz. The ground electrode was located at AFz. The electrooculogram (EOG) was recorded via electrodes located at the outer canthus of each eye and above and below participants’ left eye. The EEG was amplified using a BrainAmp DC amplifier (Brain Products GmbH, Gilching, Germany) with an initial band-pass filter of DC – 250 Hz and a sampling rate of 1000 Hz. Electrode impedances were kept below 10kΩ. EEG was also recorded during two minutes of eyes-open and two minutes of eyes-closed resting-state periods immediately before the learning task and after the judgement task.

EEG analysis was performed in MATLAB 2017b (The MathWorks, Natick, USA) using custom scripts in conjunction with the Fieldtrip toolbox (Oostenveld et al., 2011). EEG data were re-referenced offline to the average of both mastoids and band-pass filtered from 1 – 40 Hz using a two-pass Butterworth IIR filter (implemented in ft_preprocessing). Data were then epoched from -200ms to 13s relative to the onset of each picture-sentence pair for both fixed and flexible sentences, and corrected for ocular artefacts using Infomax Independent Component Analysis (Bell & Sejnowski, 1995; implemented in runica.m). Components demonstrating clear EOG artefacts were removed (median components rejected = 3, range = 2 – 6) and electrodes showing strong artefacts were visually inspected and subsequently interpolated with surrounding electrodes based on spherical spline interpolation (total channels interpolated *n* = 2; Perrin et al., 1989).

### EEG data analysis

The aim of the analysis was to characterise the oscillatory and aperiodic dynamics underlying the initial encoding of complex grammatical rules. To this end, we computed differences between fixed and flexible word order sentences in the following five spectral features of the EEG recorded during the learning phase: mean power density within individualised delta, theta, alpha, and beta bands, and the (inverse) slope of the 1/*f* spectral distribution (i.e., power-law exponent). We then used these metrics to investigate whether trial-level variation in oscillatory power and 1/*f* slope during the learning task predicted behavioural performance on the judgement task. We also tested whether interactions between oscillatory and aperiodic activity afford unique information predicting behavioural performance.

#### Spectral decomposition and power-law exponent χ estimation

The power-law scaling exponent χ, which summarises the rate of decay of the power spectrum in double-logarithmic co-ordinates, was estimated using the Irregular-Resampling Auto-Spectral Analysis toolbox (IRASA v1.0; Wen & Liu, 2016). Briefly, this technique seeks to separate oscillatory from aperiodic (random fractal) components by iteratively resampling the power spectrum across a range of non-integer factors *h* and their reciprocals 1/*h* (here, *h* = 1.1 to 1.95 in steps of 0.05). This procedure shifts any narrowband components away from their original location along the frequency spectrum while leaving the distribution of the fractal component intact. The median of the resampled spectral estimates is then calculated in order to strip the spectrum of narrowband peaks. For a more detailed treatment of the IRASA method, see Wen and Liu (2016).

Trial data from each EEG channel were divided into two non-overlapping 4500ms segments corresponding to the picture and sentence presentation phases, respectively. Picture segments were timelocked to 500ms post-stimulus onset; sentence segments were timelocked to 100 ms prior to the first word onset. Both segments were further subdivided into seven 1800ms epochs (25% overlap) and separately passed to amri_sig_fractal.m for spectral parameter estimation. Once the fractal component had been recovered from each power spectrum, it was parameterised using amri_sig_plawfit.m. This function rescales the frequency spectrum to achieve equally-spaced intervals in log-space before fitting a linear regression to a subregion of the double-log transformed fractal spectrum (here, ∼1.9 – 15.8 Hz, corresponding to an evaluated frequency range of 1 – 35 Hz; see Gerster et al. 2022, for further details). The absolute value of the regression slope coefficient was taken as the χ exponent.

To ensure the robustness of our analysis, we compared our estimates of the χ exponent against those derived using the more recently-developed ‘FOOOF’ method (Donoghue et al., 2020). Briefly, this technique attempts to separate narrowband oscillatory peak components from broadband aperiodic activity by iteratively fitting Gaussian functions to the spectrum, and deleting these components until no further deviations from background activity can be detected (given a predefined noise threshold). The χ exponent is then estimated by fitting a regression to the residual spectrum in double-log space, similar to the IRASA procedure (see Donoghue et al., 2020, for details).

Since FOOOF requires PSD estimates (rather than timeseries data) as its input, power spectra were derived from each epoch using the pwelch.m implementation of Welch’s (1967) modified periodogram method. Epochs were Hann-tapered and zero-padded to 2048 points to facilitate comparability with IRASA-generated spectral estimates. FOOOF was implemented via the MATLAB wrapper (v1.0.0) using the following parameter settings: peak width limits = 1 – 12 Hz, maximum number of peaks = infinite, minimum peak height = 0, peak threshold = 2 S.D., aperiodic mode = fixed, evaluated frequency range = 1 – 35 Hz.

#### Spectral band power estimation

In order to quantify narrowband changes in spectral power independent of underlying changes in aperiodic activity, mean power densities were estimated following the subtraction of the mean regression fit of the aperiodic component from the PSD (spectra averaged across epochs within each segment). This residual, ‘oscillatory’ spectrum was half-wave rectified (negative values set to zero) and divided into the four frequency bands of interest. Notably, the limits of each frequency band were adapted for each participant on the basis of their resting-state EEG. Specifically, the boundaries of each frequency band were calculated according to the harmonic frequency architecture proposed by Klimesch (2012; 2013; and which is in line with previous work, e.g., Corcoran et al., 2018, Doppelmayr et al., 1998, Sauppe et al., 2021), in which the centre frequency of each successive band constitutes a harmonic series scaled in relation to the individual alpha frequency (IAF). To avoid the potential overlap of neighbouring frequency bands, we determined lower and upper frequency bounds using the following formulae:

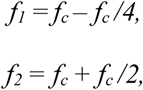

where *f*_c_ is the centre frequency (based on the IAF-scaled harmonic series), *f*_*1*_ the lower bound, and *f*_*2*_ the higher bound of a given frequency band.

IAF estimates used to determine *f*_*c*_ were obtained from a set of parieto-occipital electrodes (P3/P4/O1/O2/P7/P8/Pz/Iz) using the *restingIAF* package (v1.0.3; Corcoran et al., 2019; see also Cross et al. 2020b). This method applies a Savitzky-Golay filter (frame width = 11 bins, polynomial order = 5) to smooth and differentiate the power spectrum prior to estimating the peak frequency within a specified frequency range (here, 7—14 Hz). Peak estimates were averaged across channels, with a minimum of 3 channel estimates required to return an IAF for a given recording. Estimates derived from pre- and post-session eyes-closed resting states were then averaged for each participant using meanIAF.m. For further details on this algorithm, see Corcoran and colleagues (2018).

Having determined IAF-anchored bounds for the delta, theta, alpha, and beta bands, power within each band was quantified using the mean power density metric proposed by Westfall (1990):

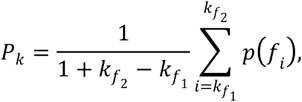

where *p(f*_*i*_) is the power estimate of the *i*^th^ frequency bin, and *f*_1_ and *f*_2_ index the lower and upper bounds of the individualised frequency band *k*, respectively. An advantage of this approach is that power estimates are scaled by spectral range, thus controlling for differing frequency bandwidths both within and between individuals.

### Statistical analysis

We used *R* v.4.0.0 (R Core Team, 2020) and the packages *lme4* v.1.1.27.1 (Bates et al., 2015), *lmerTest* v.3.1.2 (Kuznetsova et al., 2017), *ggeffects* v.4.1.4 (Lüdecke, 2018), *car* v.3.0.7 (Fox et al., 2011), *tidyverse* v.1.3.0 (Wickham et al., 2019), *mgcv* v.1.8.36 (Wood, 2006), *mgcViz* v.0.1.9 (Fasiolo et al., 2019), *rgl* v.0.1.54 (Nenadic & Greenacre, 2007), *ggpubr* v.0.4.0 (Kassambara (2020), *cowplot* v.1.0.0 (Wilke, 2019), and *eegUtils* v.0.7.0 (Craddock, 2022). For linear models, contrasts for categorical variables were sum-to-zero contrast coded, with coefficients reflecting deviation from the grand mean (Schad et al., 2020).

#### Generalised additive mixed models

Generalized additive models (GAMs) are a nonparametric extension of the standard linear regression model that substitute a linear predictor variable *x* with a smooth function *f(x)* (Hastie & Tibshirani, 1987, 1990; Wood, 2017). Generalized additive mixed models (GAMMs; Lin & Zhang, 1999) constitute a further extension that incorporates random effects components within the GAM framework (Pedersen et al., 2019). Together, these innovations offer an elegant solution to the problem of autocorrelation amongst residuals induced by (1) attempting to fit linear models to non-linear relationships, and (2) non-independence (or nesting) of observations (e.g., repeated measures within participants or items; Baayen et al., 2008).

Here, GAMMs were constructed to investigate how the exponent χ, and the mean power density *P* for each *k*^*th*^ frequency band (delta, theta, alpha, and beta), fluctuate during artificial grammar learning. Trial-level χ and *P*_*k*_ estimates from the sentence processing phase of each trial were modelled as a function of learning time (trial number), sensor space (2D Cartesian co-ordinates), and sentence type (fixed, flexible). Estimates from the preceding image presentation phase were treated as a baseline measure of spectral activity. Random factor smooth interactions were included to account for individual differences in the functional relationship between spectral features and time-on-task (see Baayen et al., 2017, Corcoran, Macefield, & Hohwy, 2021, for similar approaches). Each GAMM took the following general form:

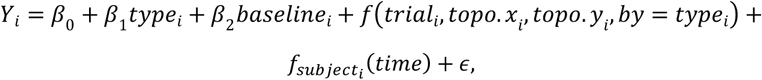

where *Y*_*i*_ is the *i*^th^ observation of spectral feature *Y, β*_0_ is the model intercept, *β*_1_*type* is a factor encoding the main effect of Sentence Type, *β*_2_*baseline* is a covariate encoding the corresponding observation for *Y* during the baseline period, *f(*., by = *type)* is the tensor product interaction between the learning time (*trial*) and sensor space (*topo*.*x, topo*.*y*) covariates for each level of Sentence Type, *f*_*subject*_ is the by-participant factor smooth for time-on-task, and *ε* is a *t* distributed error term (since response variables were heavy-tailed). Note that marginal smooths for sensor space co-ordinates were treated as isotropic (i.e., assumed to share a common scale).

GAMMs were estimated using the bam() function of the *R* package mgcv (Wood, 2011). Models were fit using the Fast REML method. *P*_*k*_ estimates for both the baseline and sentence processing period were log_10_ transformed prior to model inclusion. Models were fit with tensor product interaction smooths in order to enable ANOVA-decomposition of main effect and interaction components (Wood, Scheipl, & Faraway, 2013). All tensor product smooths were fit using low rank thin plate regression splines as their basis function (Wood, 2003, 2017). Factor smooths were fit with a first-derivative penalty in order to shrink participant-level smooths towards the population-level. An additional shrinkage penalty was imposed on the smoothing penalty null space to enable automated model reduction (see Marra & Wood, 2011). Type was entered as an ordered factor with Fixed assigned as the reference level, hence model terms involving a Sentence Type interaction assess the difference between Fixed and Flexible condition splines (see van Rij et al., 2016).

#### Linear mixed-effects models

The relationship between aperiodic and oscillatory power during grammar learning with behavioural performance on the judgement task was assessed using linear mixed-effects models. Behavioural performance was operationalised using the discrimination index (d’). d’ is defined as the difference between the z transformed probabilities of hit rate (HR) and false alarm rate (FA; i.e., d’=z[HR] – z[FA]). These models took the following general form:

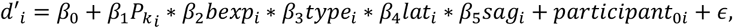

where *P*_*k*_ is mean baseline-corrected (i.e., sentence presentation – pre-sentence interval) power density in the frequency band of interest (i.e., delta, theta, alpha, beta), *bexp* is the baseline-corrected (i.e., sentence presentation – pre-sentence interval) exponent of the aperiodic 1/*f* slope, *type* refers to Sentence Type (fixed, flexible), *sag* is Sagittality (anterior, central, posterior) and *lat* refers to Laterality (left, midline, right). Participant ID (*participant*) was modelled as a random intercept. ε refers to a Gaussian-distributed error term.

Type II Wald χ^2^-tests from the *car* package (Fox et al., 2011) were used to provide *p*-values. An 83% confidence interval (CI) threshold was adopted for visualisations, which corresponds to the 5% significance level with non-overlapping estimates (Austin & Hux, 2002; MacGregor-Fors & Payton, 2013). General linear models were performed to assess the relationship between baseline corrected oscillatory power and aperiodic 1/*f* slope between fixed and flexible word orders. Baseline-corrected oscillatory power values were log_10_ transformed prior to model inclusion. All data, as well as analysis scripts (MATLAB and R) are available on the OSF platform: https://osf.io/7yr46/; Cross, Corcoran, Schlesewsky, Kohler, &. Bornkessel-Schlesewsky, 2022). For a schematic visualisation of EEG signal processing and statistical analysis steps, see Figure 2.

**Figure 2.**
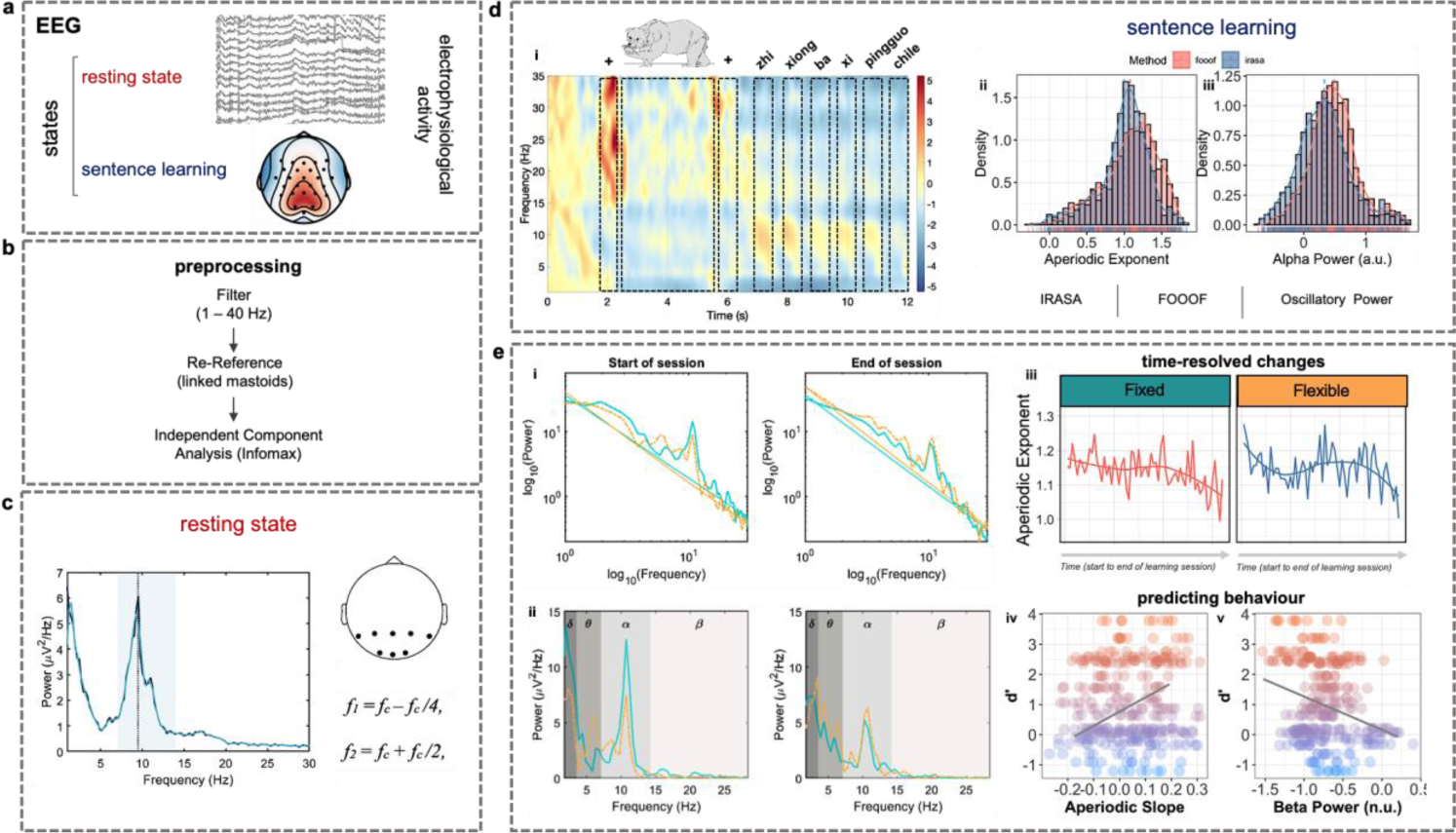
Schematic of the EEG recording, pre-processing, signal, and statistical analysis procedures. **a.** Neurophysiological signals were recorded at rest and during the sentence learning task using a 32-channel EEG system. **b**. The EEG signal was filtered, re-referenced and subjected to an independent component analysis. **c**. The individual alpha frequency (IAF) was estimated per participant from resting-state EEG recordings based on an occipital-parietal electrode cluster (see topoplot). Peak frequencies within the alpha band (7-14 Hz; light blue shading) were identified using restingIAF, an automated procedure that smoothes and differentiates the power spectrum before estimating the average IAF (dotted line) across selected channels. IAF estimates were subsequently used to calculate participant-specific delta, theta, alpha and beta centre frequencies (f_c_) and bandwidths (f_1_, f_2_) for the time-frequency decomposition of the sentence learning task. **d**. (i) Grand-average time-frequency representation of fixed word order sentences during the learning session. Dashed black boxes correspond to the presentation of elements in the stimulus train above. (ii) Histograms illustrating the distribution of the aperiodic exponent and alpha power estimated using FOOOF (pink) and IRASA (blue). **e**. (i) Single-subject power spectral density (PSD) plots at the beginning (left) and end (right) of the sentence learning task. Straight turquoise and tan lines represent the IRASA-based aperiodic regression fit for fixed and flexible word order sentences, respectively. (ii) PSD plots illustrating power in the IAF-derived delta (*δ*), theta (*θ*), alpha (*α*) and beta (*β*) bands after subtraction of the aperiodic regression fit depicted in (i). (iii) Raw data at electrode Cz illustrating the analysis performed to estimate time-varying modulations in the aperiodic exponent between fixed and flexible word orders. (iv) scatterplots illustrating the analyses predicting behaviour (i.e., judgement accuracy [d’]) from the sentence judgement task from aperiodic and oscillatory activity derived from the sentence learning task.

## Results

### Task performance

The results on the judgement task are visualised in Figure 3. Participants performed moderately well on the judgement task, with a mean d’ score of 1.02 (range: -1.20 – 3.78) and mean reaction time of 878.08 ms (range: 254.75 – 2076.83). There is clearly a high degree of inter-individual variability across both fixed and flexible sentences; however, flexible sentences had greater variability in d’ scores, while fixed grammatical sentences had faster responses overall. For a detailed report and interpretation of these behavioural data, see Cross et al. (2020a).

**Figure 3.**
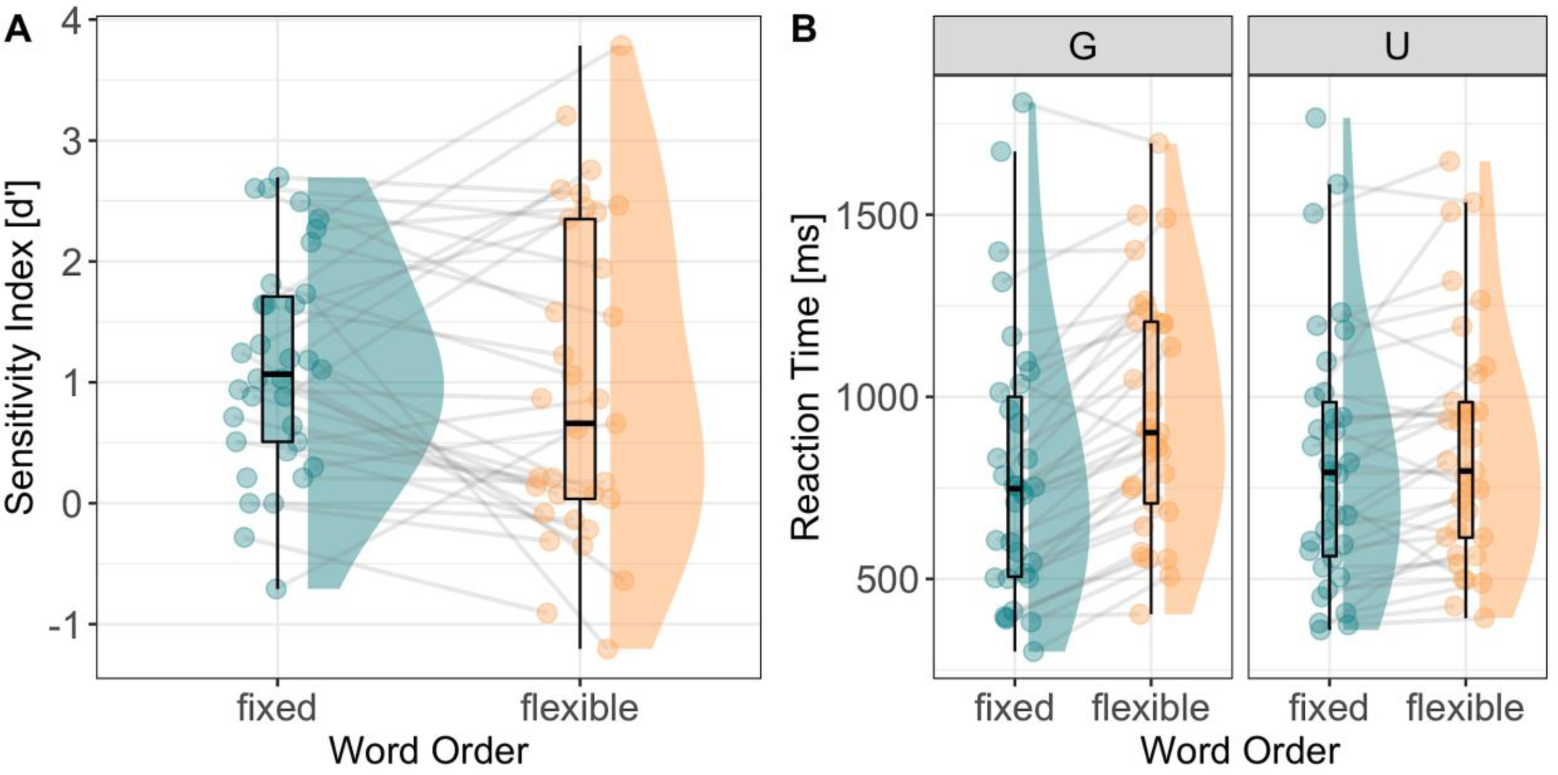
Raincloud plots illustrating the behavioural responses during the sentence judgement task. (A) Mean d’ scores (x-axis) for Fixed and Flexible sentence types. (B) Mean reaction time (ms; x-axis) for Grammatical (left) and Ungrammatical (right) Fixed and Flexible sentence types. Individual data points represent the mean for each participant, while the lines join within-participant differences between fixed and flexible word order sentences.

### Neurophysiological results

Individual alpha frequency estimates varied between participants (M_IAF_ = 9.78, *SD* = 0.96), resulting in a range of participant-specific frequency bands (summarised in Table 1). A full list of participant-specific IAFs and frequency bandwidth are available on the OSF repository.

**Table 1.** Mean lower (*f*_*1*_) and upper (*f*_*2*_) frequency bounds for the delta, theta, alpha and beta bands. Participant-specific range provides the lowest and highest frequency band limits based on single-participant estimates, as is also provided for IAF estimates.

| Band | Mean $f_1$ (SD) | Mean $f_2$ (SD) | Participant-Specific Range |
| --- | --- | --- | --- |
| Delta | 1.83 (0.18) | 3.67 (0.36) | 1.42 – 4.33 |
| Theta | 3.67 (0.36) | 7.34 (0.72) | 2.83 – 8.66 |
| Alpha | 7.34 (0.72) | 14.7 (1.45) | 5.67 – 17.33 |
| Beta | 14.7 (1.45) | 29.4 (2.90) | 11.36 – 34.67 |
| IAF | -- | -- | 7.57 – 11.55 |

*Note.* SD = standard deviation; IAF = individual alpha frequency. Participant-specific range provides the absolute lowest and upper band limits.

#### Aperiodic and oscillatory changes across time and space during language learning

Neurophysiological signals are non-stationary, showing dynamic changes over time as a function of endogenous and exogenous factors (e.g., Donoghue, Schaworonkow & Voytek, 2021), such as attentional fluctuations and the complexity of incoming sensory information (Waschke et al., 2021). However, neurophysiological signals are typically analysed using linear models, which often do not capture non-linear modulations in neural activity, particularly over time. Here, we examine how aperiodic and oscillatory dynamics evolve over time during language learning, focusing specifically on the way in which spectral activity varies across sentence types (fixed vs flexible word orders). Estimated changes in aperiodic and oscillatory spectral activity across learning task conditions are illustrated in Figure 5 (for topographical maps, see Figure 6).

Comparisons between IRASA and FOOOF showed that FOOOF provided higher exponent estimates than IRASA, irrespective of sentence type (fixed, flexible; Figure 4) and were also more variable (M_FOOOF_ = 1.08, *SD* = 0.48; M_IRASA_ = 1.00, *SD* = 0.42). However, exponent estimates between IRASA and FOOOF were highly positively correlated across both fixed (ρ = 0.75, *p* < .001, 83% CI = [.63, .84] and flexible (ρ = 0.76, *p* < .001, 83% CI = [.63, .85]; Figure 4B) word order sentences. These observations were complemented by a linear mixed-effects regression which revealed that while FOOOF had overall higher exponent estimates than IRASA (*β* = 0.02, *se* = 0.004, *p* < .001), exponent estimates between FOOOF and IRASA did not vary by sentence type (*β* = 0.0005, *se* = 0.004, *p* = .89; Figure 4C). These observations are consistent with simulations reported by Donoghue et al., 2020; however, given that there was no significant interaction between method (FOOOF, IRASA) and sentence type (fixed, flexible), we present the IRASA-based analysis.

**Figure 4.**
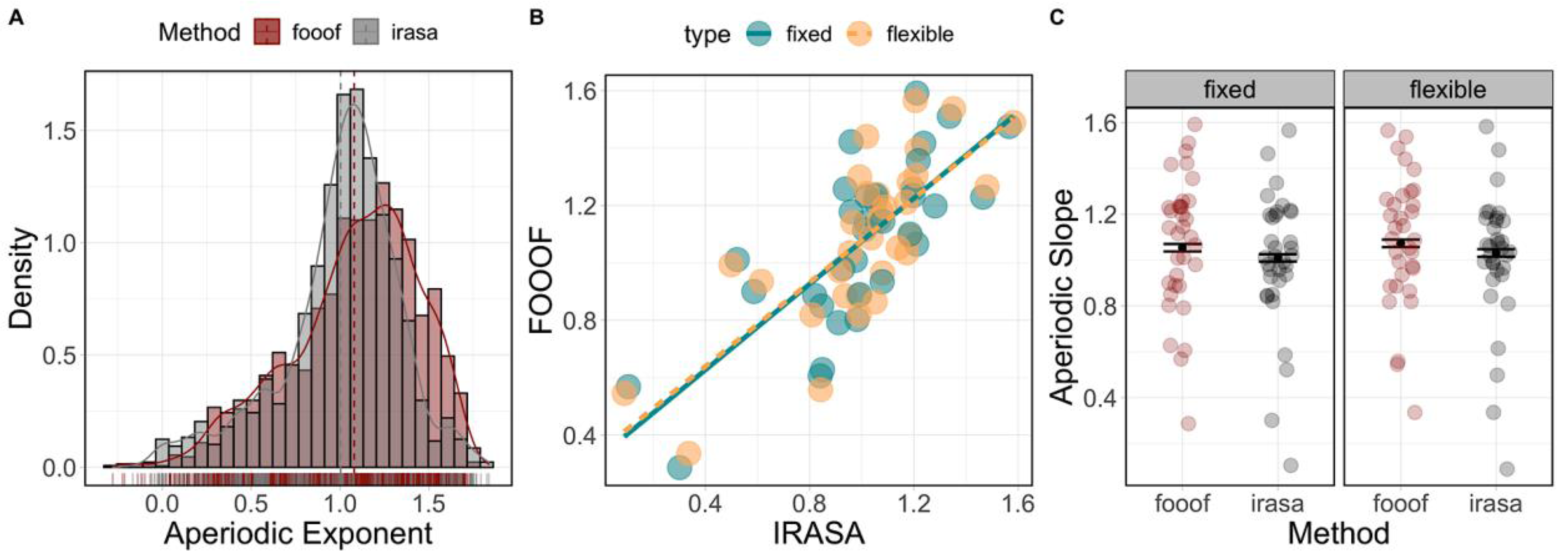
Comparison between FOOOF and IRASA exponent estimates. (A) Histogram illustrating the distribution of exponent estimates derived from FOOOF (red) and IRASA (grey). (B) Scatterplot showing the relationship between FOOOF (y-axis) and IRASA (x-axis) between fixed (turquoise) and flexible (tan) sentences. (C) Relationship between the aperiodic exponent (y-axis; higher values indicate a steeper exponent), method (x-axis; FOOOF, IRASA), and sentence type (left facet = fixed, right facet = flexible). Bars represent the 83% confidence interval around group-level expected marginal mean estimates. Dots represent individual data points per participant for aggregated data.

**Figure 5.**
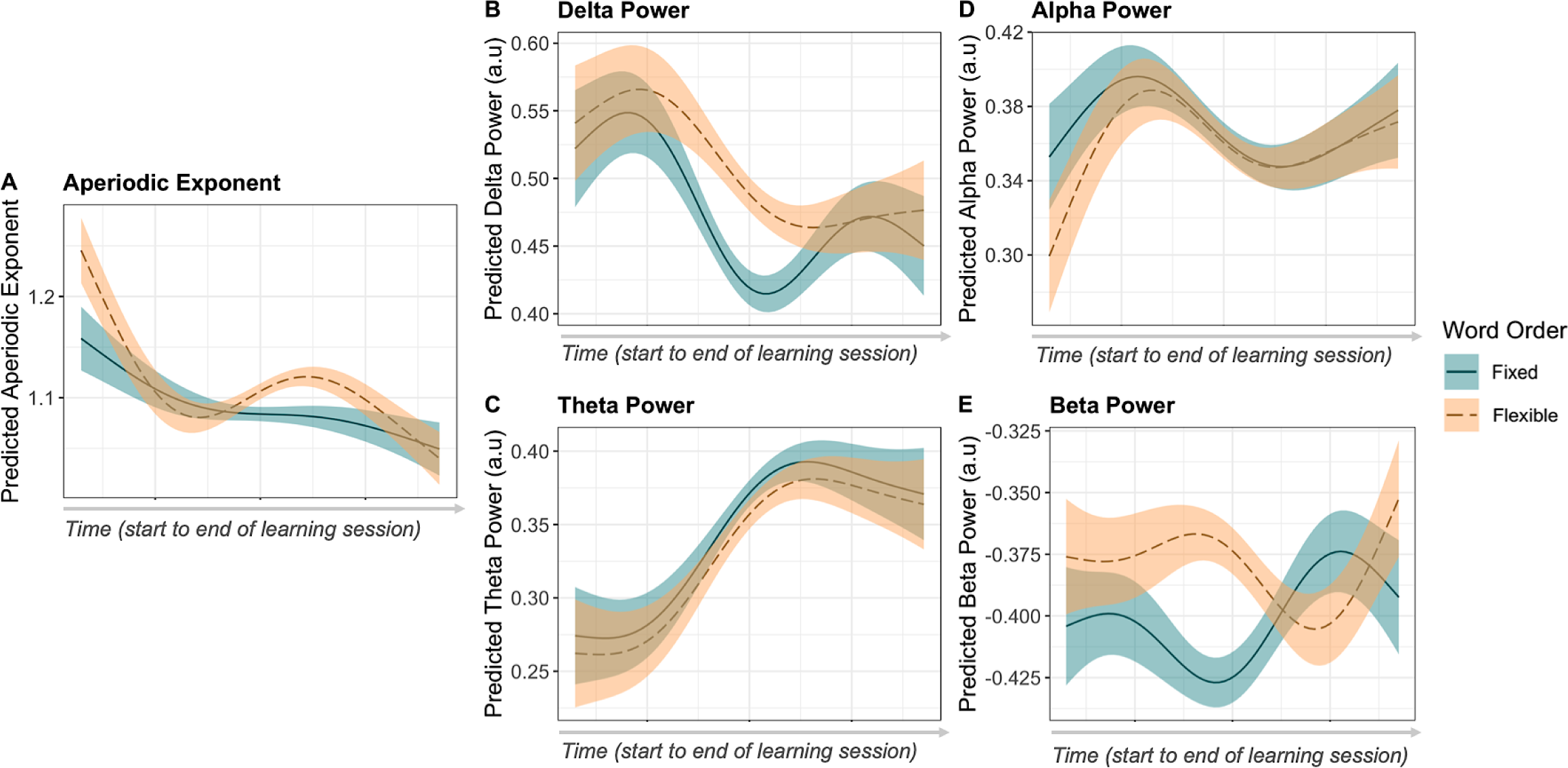
Modelled effects for changes in the aperiodic exponent (A), delta (B), theta (C), alpha (D) and beta (E) activity across the learning task for fixed (solid line) and flexible (dashed line) word order rules. Time from beginning to the end of the learning task is represented on the x-axis.

The χ-exponent GAMM revealed the 1/*f* slope was steeper on average for Flexible compared to Fixed word order sentences (*β* = 0.02, *SE* = 0.008, *F*(1) = 5.84, *p* = .015). Visualisation of smooth terms (Figure 5A) revealed that exponent values tended to decrease (indicating a flattening of the 1/*f* slope) over the course of the learning period; however, Flexible trials evoked higher values (steeper 1/*f* slopes) at the beginning and during the second half of the session, relative to Fixed trials (Trial × Sentence Type estimated degrees of freedom [*edf*] = 3.87, *F* = 29.67, *p* < .001). This model further revealed significant topographic differences between conditions, with Flexible word orders evoking higher exponent values over fronto-central regions compared to Fixed word order sentences by the end of the session (Trial × Sagittality × Sentence Type *edf* = 1.78, *F* = 1.13, *p* < .001; see Figure 6; for full summary tables of all models, see Appendix).

Mean delta power was higher on average during Flexible compared to Fixed word order trials, although this difference was not significant (*β* = 0.03, *SE* = 0.016, *F* = 3.68, *p* = .055). However, visualisation of smooth terms (Figure 5B) revealed a complex pattern whereby delta power increased over early trials, followed by a marked decrease that was more pronounced in response to Fixed than Flexible sentence stimuli (*edf* = 2.74, *F =* 9.66, *p* < .001). This interaction was significantly modulated by sagittality, with between-condition differences (Fixed versus Flexible) in mean power density increasing over fronto-central electrodes as a function of time (*edf* = 2.68, *F* = 1.54, *p* < .001). Theta power was non-significantly lower on average during Flexible compared to Fixed rule learning (*β* = - 0.01, *SE* = 0.014, *F* = 0.60, *p* = .437). Again, smooth terms revealed a nonlinear pattern of spectral fluctuation, whereby theta power evinced a sigmoidal shape over the course of rule-learning (Figure 5C). This pattern was similar across both conditions, with Fixed sentence stimuli tending to evoke increased theta power (*edf* = 0.824, *F* = 1.25, *p* = .014). This pattern of activity varied as a function of topography, with the difference between conditions being more accentuated across lateralised and posterior sites, as illustrated in Figure 6 (*edf* = 7.63, *F* = 0.22, *p* = .003).

Alpha power tended to increase over the early and later trials of the learning task, although this pattern was interrupted by a marked decline during the middle of the session (*edf* = 3.20, *F* = 12.50, *p* < .001). Flexible word orders evoked less alpha power than Fixed word orders at the beginning of the session, but was similar thereafter (*edf* = 3.20, *F* = 13.46, *p* < .001; Figure 5D). This difference was most pronounced over left-lateralised and frontal sites (*edf* = 2.27, *F* = 0.35, *p* = .012). Finally, the beta power model revealed significant differences in the nonlinear profile of power dynamics across the learning session. In fact, Fixed and Flexible trials evoked markedly different patterns of activity: beta power showed an approximately triphasic response to Fixed sentence stimuli that was mirrored by the response to Flexible stimuli (*edf* = 2.94, *F* = 30.08, *p* < .001; Figure 5E). The strongest beta response was observed over frontal and temporal regions (*edf* = 16.59, *F* = 33.04, *p* < .001), particularly toward the beginning of the learning phase for Flexible word order sentences (*edf* = 6.82, *F* = 1.24, *p* < .001).

Taken together, these data illustrate dynamic changes in both aperiodic and oscillatory activity as a function of different word order rules during learning. Both the aperiodic slope and delta power tended to decrease over time, while theta power tended to increase. By contrast, alpha and especially beta power evinced more complex dynamics as participants learnt different word order rules.

#### Task-related aperiodic and oscillatory activity are dynamically related during language learning

Neurophysiological signals are dominated by transient oscillatory and broadband aperiodic activity; however, in the study of the oscillatory correlates of higher-order language processing, aperiodic activity is rarely considered, with little known regarding its influence on task-related oscillatory activity (cf. Cross et al., 2021). Here, we examined the associations between task-related oscillations in individualised (i.e., anchored on participants’ IAF) frequency bands and aperiodic activity during the learning of different word order rules. There was non-significant positive association between delta power and the aperiodic slope (*β* = 0.58, *p* = .05, *R*^*2*^ = 0.02). There was no relationship between theta power and the aperiodic slope (*β* = 0.09, *p* = .77, *R*^*2*^ = -0.04); however, there was a significant negative association between alpha power and the aperiodic slope (*β* = -2.88, *p* < .001, *R*^*2*^ = 0.33), which did not vary by sentence type. Finally, there was no significant relationship between task-evoked beta power and the aperiodic slope (*β* = 0.31, *p* = .49, *R*^*2*^ = -0.04; for a visualisation of these associations, see Figure 7). These results indicate that aperiodic and narrowband spectral estimates may afford complementary information about learning and task performance. Based on this, we now examine whether such aperiodic and (putative) oscillatory activity interact to predict performance on the sentence judgement task.

**Figure 6.**
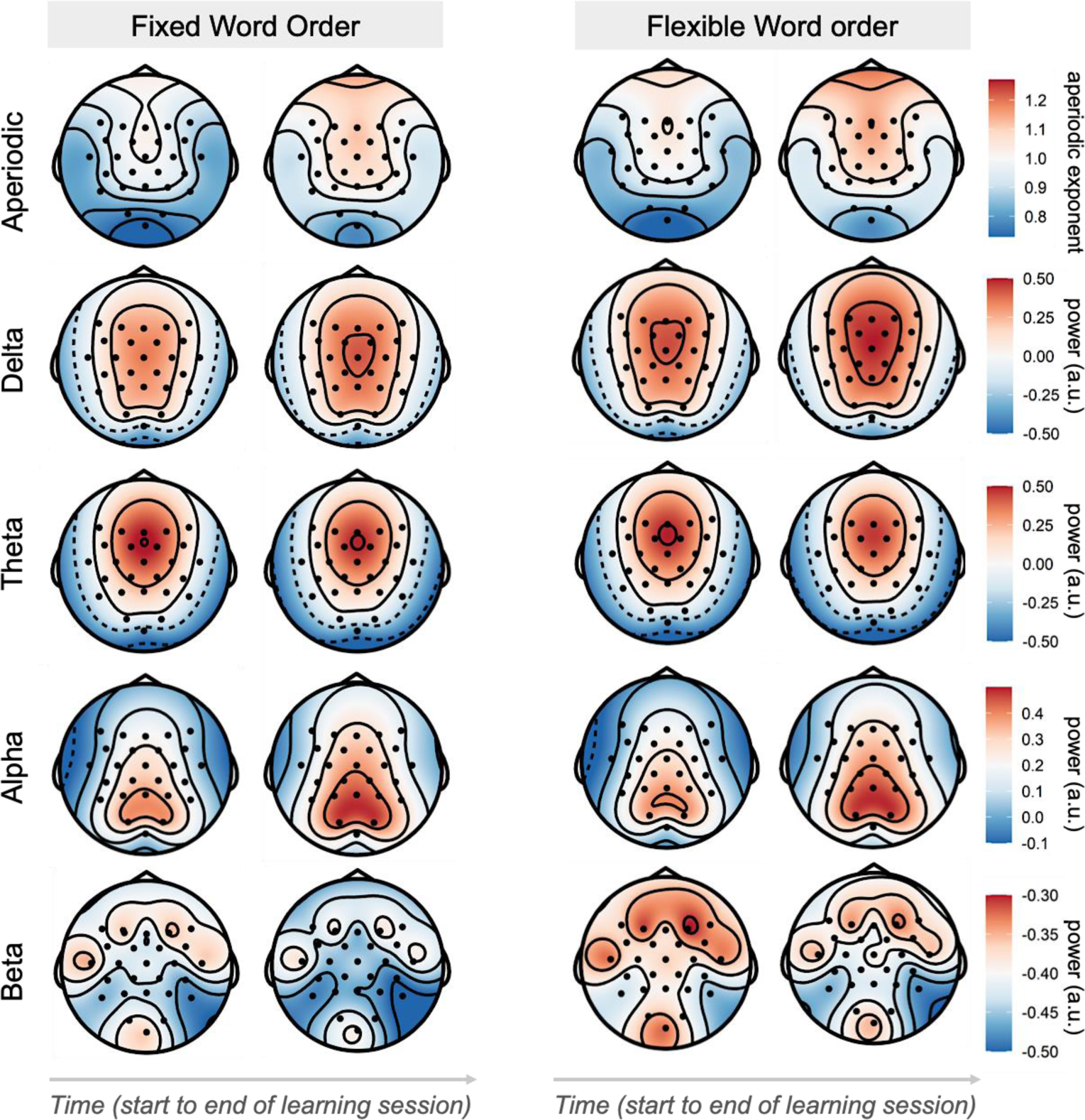
Difference in topographical distribution of aperiodic and oscillatory activity between fixed and flexible word order sentences at the beginning and end of the sentence learning task.

**Figure 7.**
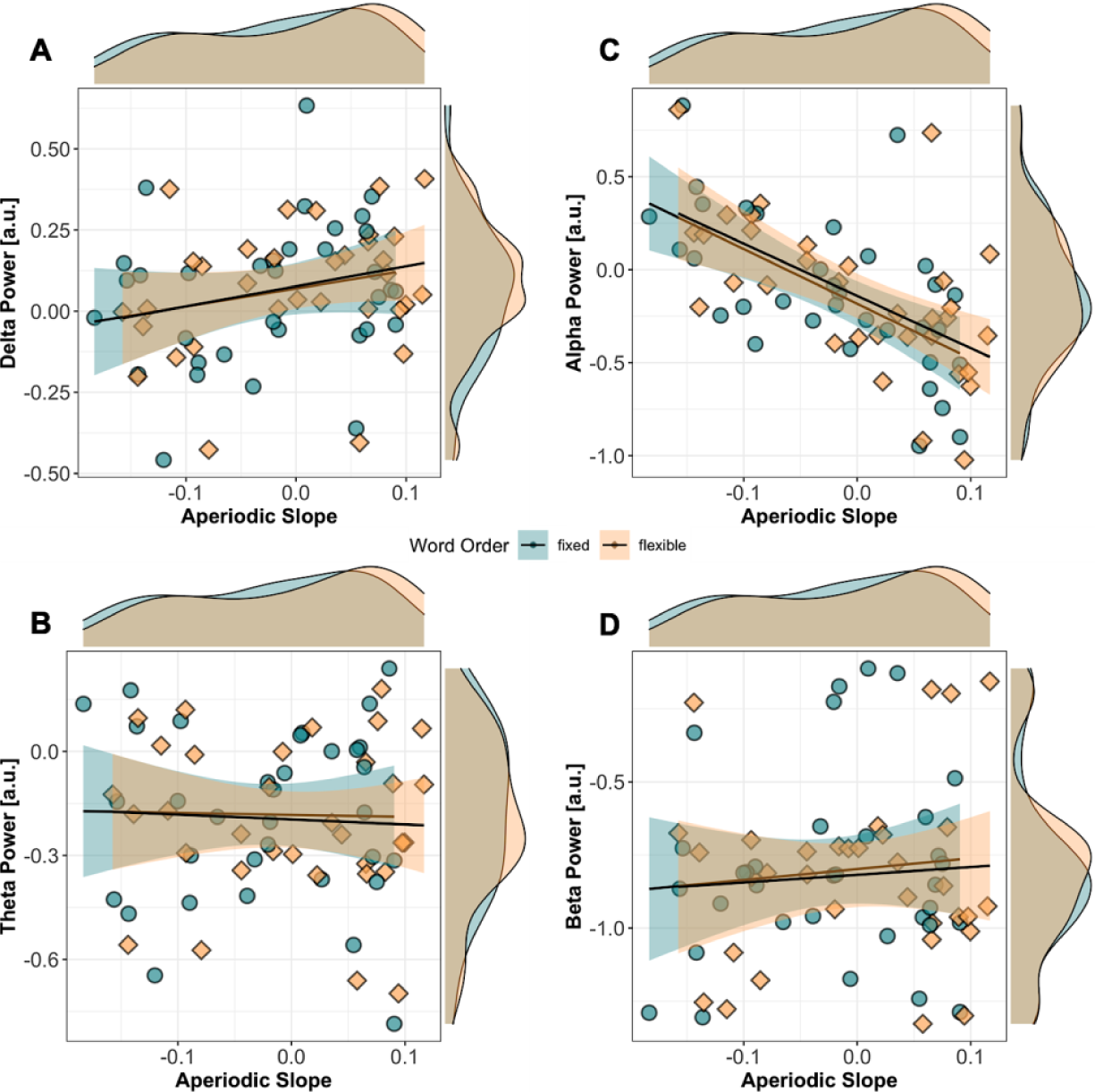
Association between task-related aperiodic slope and oscillatory power in the delta (A), theta (B), alpha (C) and beta (D) bands during the sentence learning task averaged across all channels. The aperiodic slope is represented on the x-axis (higher values indicate a steeper slope relative to the pre-sentence interval), while oscillatory power is represented on the y-axis (higher values indicate higher power relative to the pre-sentence interval). The density of observations for frequency band power and the aperiodic slope are indicated on the margins of each plot, while the fixed and flexible word order sentences are coded in turquoise and tan, respectively.

### Interactions between oscillatory and aperiodic activity modulate behavioural performance

Given the association between task-related oscillatory and aperiodic activity during the learning of complex linguistic rules, we now examine whether the 1/*f* slope and oscillatory activity interact during learning to influence behavioural performance on the sentence judgement task. For all analyses, we used linear mixed-effects regression models (for full summary tables for all models, see Appendix). For the delta model, there was a significant Power × 1/*f* Slope × Sentence Type interaction (χ^2^(1) = 12.47, *p* < .001). As shown in Figure 8A, when the 1/*f* slope was steepest and delta power was low, d’ was higher for flexible relative to fixed word orders. By contrast, when the 1/*f* slope was steep and delta power was high, performance for fixed word orders increased. For the theta model, there was a significant Power × Sentence Type (χ^2^(1) = 4.57, *p* = .03) and 1/*f* Slope × Sentence Type interaction (χ^2^(1) = 17.43, *p* < .001). Here, when the 1/*f* slope was steep, d’ scores increased for flexible sentences (Figure 8B; left). By contrast, when theta power increased, performance for both fixed sentences was higher (Figure 8B; right).

**Figure 8.**
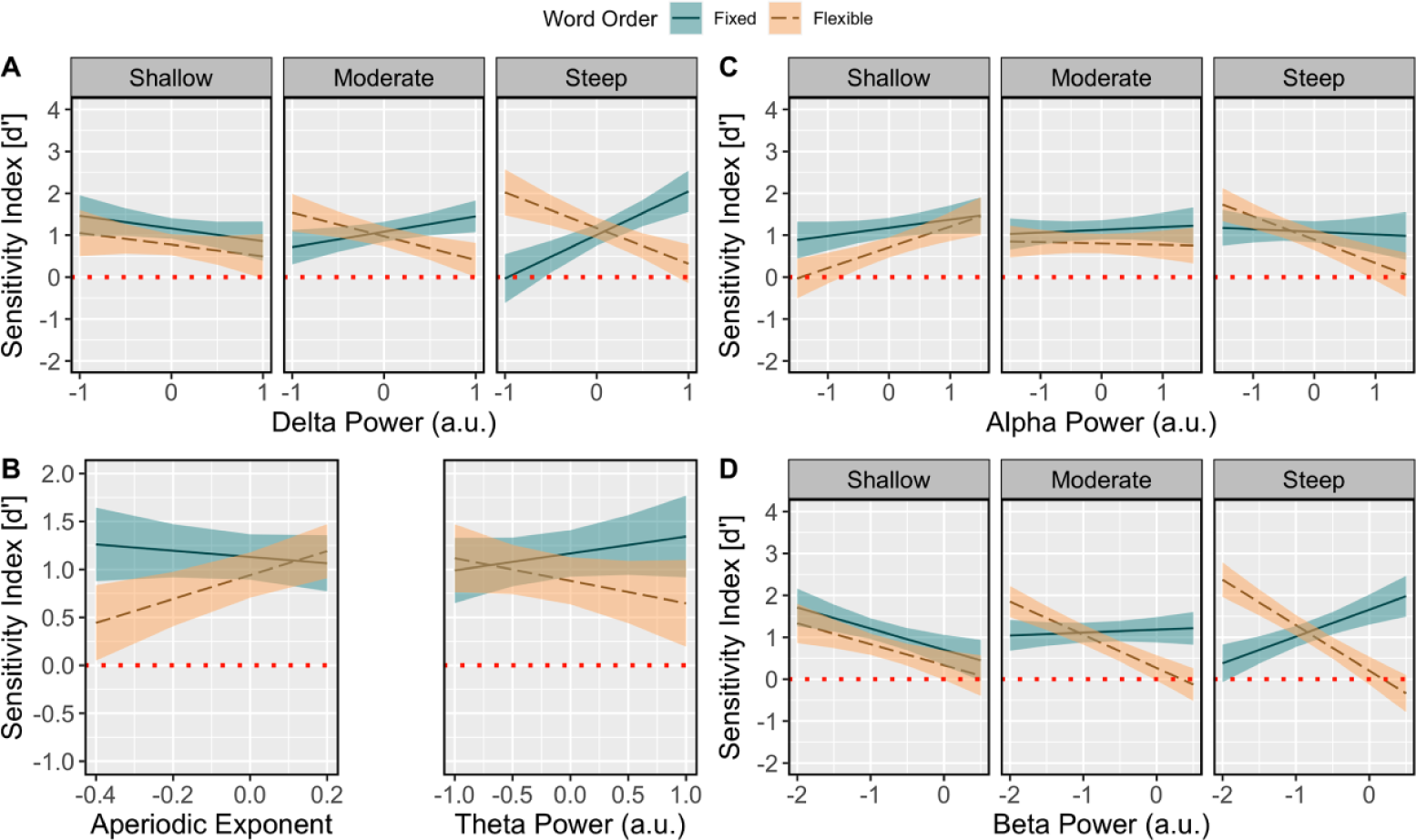
Visualisation of the relation between behavioural performance, aperiodic slope, and oscillatory delta (A), theta (B), alpha (C), and beta (D) activity. Modelled effects of task-related oscillatory activity (x-axis; higher values indicate greater power) on d’ scores (y-axis; higher values indicate better performance) for fixed and flexible word order sentences (fixed = solid line; flexible = dashed line). Task-related aperiodic 1/*f* slope estimates are faceted from shallow (left), to moderate (middle), to steep (right). Note that the trichotomisation of the aperiodic slope into shallow, moderate and steep facets is for visualisation purposes only, with the aperiodic slope being entered into all models as a continuous predictor. Note that (B) illustrates the two-way modelled interaction effects of task-related aperiodic slope (left; x-axis, higher values indicate a steeper slope) and theta power (right; x-axis, higher values indicate greater power) for fixed and flexible word order sentences (fixed = solid line; flexible = dashed line). The red dashed line indicates chance-level performance, while the shaded regions indicate the 83% confidence interval. For A, B, C and E, the x-axis reflects scaled single-trial oscillatory power estimates, with negative values reflecting a decrease in power and positive values reflecting an increase in power.

For the alpha model, there was a significant three-way Power × 1/*f* Slope × Sentence Type interaction (χ^2^(1) = 15.94, *p* < .001). As depicted in Figure 8C, when the 1/*f* slope was shallow, and as alpha power increased, d’ scores were higher for both fixed and flexible word orders. By contrast, when the 1/*f* slope was steep, and as alpha power decreased, d’ scores were lower for flexible word order sentences. Similarly, the beta model yielded a significant Power × 1/*f* Slope × Sentence Type interaction (χ^2^(1) = 30.96, *p* < .001). When the 1/*f* slope was shallow, and as beta power decreased, d’ scores were higher for both fixed and flexible word orders (Figure 8D). By contrast, when the 1/*f* slope was steep, increased beta power predicted higher d’ scores for fixed but lower d’ scores for flexible word order sentences, respectively.

Together, these results suggest that when there is a steeper 1/*f* slope, increased delta and beta power were associated with better behavioural performance, and thus better learning outcomes for fixed relative to flexible word order sentences. Further, when the 1/*f* slope was shallow and alpha power decreased, there was a general benefit in performance for both fixed and flexible word order sentences, relative to when the 1/*f* slope was steep.

## Discussion

Here, we estimated the 1/*f* slope during artificial grammar learning to characterise the influence of dynamic alterations in aperiodic and oscillatory activity on higher-order cognition. This is the first study to examine aperiodic activity and its interaction with oscillatory power in the context of language learning, with three critical findings emerging: (1) both (putative) oscillatory and aperiodic activity dynamically change over time during complex language-related rule learning; (2) the 1/*f* slope becomes steeper during the learning of complex rules, but this effect differed depending on the type of rules being learned, and; (3) learning-related aperiodic activity interacted with oscillatory power to modulate behavioural performance for both fixed and flexible word orders. These findings speak strongly to the view that aperiodic 1/*f* dynamics should be explicitly modelled or isolated as a source of variance when analysing power spectra to ensure that any oscillatory changes are not confounded with modulations in broadband aperiodic activity (Donoghue et al., 2021).

Indeed, a considerable proportion of work examining the oscillatory correlates of higher-order language processing have not explicitly accounted for modulations in broadband aperiodic activity (e.g., Bonhage et al., 2017; Corcoran et al., 2022; Kepinska et al., 2017; Lewis et al., 2016; Mai, Minett, & Wang, 2016; Prat et al., 2016; Rossi & Prystauka, 2020; c.f., Cao et al., 2022), making it difficult to determine whether oscillatory activity parsimoniously explains behavioural outcomes. By separating oscillatory and aperiodic components, we have demonstrated that the aperiodic exponent flattens across time, while, for example, theta and alpha power increase across time throughout the language learning phase. Recent computational work has highlighted the criticality of such a separation of neural signals (e.g., Donoghue et al., 2020), given that both aperiodic and oscillatory signals vary by clinical status (Robertson et al., 2019), state of consciousness (e.g., sleep versus wake; Lendner et al., 2020), and are modulated by task demands (Waschke et al., 2021). From this perspective, language studies reporting differences in oscillatory activity (e.g., increases in theta power) between experimental conditions (e.g., grammatical vs ungrammatical sentences) without accounting for broadband activity may be confounded by changes in aperiodic dynamics (Donoghue et al., 2021).

### Aperiodic and oscillatory activity are modulated by time-on-task

The potential confounding of aperiodic and oscillatory components is further compounded by the fact that neural activity is non-stationary (Donoghue et al., 2021; Kosciessa et al., 2020; Stokes & Spaak, 2016). Here, we modelled single trial fluctuations of both aperiodic and oscillatory EEG components across the learning task, revealing fine-grained temporal dynamics underlying complex rule learning. For the aperiodic component, we observed a general flattening of the slope across time for both fixed and flexible sentences; however, the slope was steeper overall for flexible sentences. The general flattening of the aperiodic slope across time is in line with previous work reporting attentional modulations of spectral exponents (Kosciessa et al., 2021; Waschke et al., 2021). As exposure to grammar rules increased with time-on-task, participants may have become more adept at allocating attention to cues relevant for successful sentence interpretation. Increased attentional modulation in accordance with learnt rules may have been accompanied by increased excitation/inhibition ratio, which reflects an increase in high-frequency power in cortical regions involved in processing task-relevant information (Cohen & Maunsell, 2011; Harris & Thiele, 2011), thus explaining the flattening of the aperiodic slope (Kosciessa et al., 2021; Waschke et al., 2021).

The observed increases in theta and alpha power over time are also consistent with previous work on complex rule and language learning (e.g., Crivelli-Decker et al., 2018; de Diego-Balaguer, Fuentemilla, & Rodriguez-Fornells, 2011; Kepinska et al., 2017). In the few studies examining the neural oscillations involved in grammar learning (e.g., de Diego-Balaguer et al., 2011; Kepinska et al., 2017), it has been demonstrated that theta and alpha synchronisation predict learning success. Here, theta and alpha power showed a non-linear increase in power across the learning task. Theta oscillations, particularly over frontal regions when recorded with scalp-EEG, are associated with plasticity-related learning and memory processes, reflecting the encoding and generalisation of new information (Eschmann et al., 2020; Khader et al., 2010). From this perspective, the observed increase in theta power for both fixed and flexible word orders may have reflected successful memory encoding and accumulating knowledge of the underlying grammatical rules.

Beta activity also displayed complex non-linear changes for fixed and flexible word orders across the learning task. Overall, beta power was higher for flexible than fixed word orders, particularly in the second half of the learning session (Figure 5E). In the native language processing literature (Bastiaansen et al., 2010; Davidson & Indefrey, 2007; Kielar et al., 2014, 2015), beta oscillations are argued to reflect prediction-related activity, with beta power increasing in highly predictable linguistic contexts, and decreasing when grammatical violations occur (for review, see Lewis et al., 2015, 2016). However, in studies on second language learning (e.g., Lewis et al. 2016), beta power increases in response to sentences with long-distance dependencies, possibly indicating more effortful processing (Meyer et al., 2013). From this position, the observed general increase in beta power for both fixed and flexible word orders across the task may reflect the accumulation of grammatical knowledge, allowing participants to better predict underlying rules of the language. Further, the marked increase in beta power for flexible word order processing may indicate more effortful processing, given that flexible word orders contain non-adjacent elements that require integration for successful comprehension (Cross et al., 2018).

### Interactions between aperiodic and oscillatory activity predict learning

Interactions between oscillatory and aperiodic activity during the learning task also predicted subsequent behavioural performance. Increased alpha power predicted an increase in performance for fixed word orders when the aperiodic 1/*f* was shallow, while a decrease in alpha power predicted higher performance for flexible word orders when the aperiodic slope was steep. By contrast, when the aperiodic slope was shallow, a decrease in beta power (i.e., beta desynchronisation) was associated with improved behavioural performance for both fixed and flexible word orders. Further, when the aperiodic slope was steep, the relationship between beta desynchronisation and flexible word order processing was stronger, but the inverse was observed for fixed word order sentences.

The effect of differing levels of 1/*f* slope on, for instance, beta power and behavioural performance likely reflect more nuanced inter-individual differences in information processing capacities (Dziego et al., 2022; Immink, Cross et al., 2021; Thuwal, Banerjee, & Roy, 2021), which may explain behavioural gains that are otherwise related to the manifestation of oscillatory activity (e.g., Kepinska et al., 2017). For example, here we observed that a decrease in beta power predicted better behavioural performance for flexible rules, while the inverse was seen for fixed word order rules. From this perspective, a steeper slope may be more conducive for learning more complex information based on distinct neural dynamics, reflecting a decrease in the excitation/inhibition balance, and thus a decrease in high-frequency activity (Cohen & Maunsell, 2011; Harris & Thiele, 2011; Waschke et al., 2021). A reduction in high-frequency activity has been associated with error-driven learning (Luft, Takase, & Bhattacharya, 2014; Luft, 2014; Tan, Jenkinson, & Brown, 2014) and predictive processing-based activity (Bastos et al., 2012; Arnal & Giraud, 2012), particularly in the context of language comprehension (Cross et al., 2018; Lewis & Bastiaansen, 2015; Lewis et al., 2016). As such, a steeper 1/*f* slope, which was observed for flexible relative to fixed word order rules across the learning task (Figure 5A), may be foundational for task-related oscillatory activity during higher-order language learning.

The oscillatory-based findings are also broadly consistent with previous work (e.g., Kepinska et al., 2017), but reveal fine-grained patterns of spectral activity between word order variations, which may be explained by cue-integration-based models of language processing (Bates et al., 2001; Bornkessel & Schlesewsky, 2006; Bornkessel-Schlesewsky et al., 2015; Kaufeld et al., 2020; Martin, 2016). Under this framework, cues that are differentially weighted according to the probabilities of the language are integrated to comprehend incoming linguistic input (e.g., sentences). Here, fixed word orders contained linear order-based cues, which are analogous to English, while flexible word orders required animacy-based cues for interpretation. From this perspective, and in line with previous work on sequence processing (Crivelli-Decker et al., 2018; Kikuchi, et al., 2018; Wang et al., 2019), increased beta power likely reflected the propagation of top-down predictions during the learning of fixed word orders (Cross et al., 2018). In fixed sentences, the first noun is invariably the Actor, and as such, predictions are constrained to anticipating that the second noun will be the Undergoer, while also containing a verb-final construction. Therefore, due to the strong sequence dependence in fixed word orders, precision-weighted predictions would likely increase linearly across the sentence, manifesting in increased beta power (Arnal, 2012; Cross et al., 2018; Lewis & Bastiaansen, 2015).

The inverse relationship with flexible word order processing – which was predicted by a reduction in beta power– can also be explained under this framework. Given that flexible word orders contain either Actor-first or Undergoer-first constructions, predictions cannot be based on the linear position of the words, and instead must be driven by the integration of (non-adjacent) animacy-based cues to arrive at an accurate sentential percept. Given that our sample consisted of native monolingual English speakers (a language that relies heavily on word order cues; Bates et al., 2001; Bornkessel-Schlesewsky, et al. 2011; MacWhinney et al., 1984), a reduction in beta power during flexible word order processing likely reflected prediction errors and internal model updating. That is, beta desynchronization during the learning of flexible word orders may have reflected internal model updating based on mismatches with predicted and actual sensory input, while an increase in beta power during fixed word order processing likely reflected the accumulation of top-down predictions based on our sample of native English speakers’ preference for word-order-based cues. Importantly, this interpretation is consistent with temporal sequence learning paradigms, where beta power increases for fixed relative to “random” sequences (Crivelli-Decker et al., 2018), which also aligns with the observed beta power increase from the second half the learning task for fixed relative to flexible word orders (Figure 5E).

Alpha activity showed a similar interaction: when the 1/*f* slope was steep, reduced alpha power (i.e., alpha desynchronisation) predicted flexible word order processing. Alpha power reductions during language comprehension may reflect goal-directed processing and enhanced allocation of attentional resources, which is required for the successful learning of flexible word orders (Kepinska et al., 2017), given that they deviate from the canonical English word order (Bates et al., 2001). This interpretation is in line with evidence demonstrating that alpha oscillations reflect rhythmic cortical gating by alternating the activation of task-relevant cortical regions while inhibiting the processing of task-irrelevant information (Chapeton et al., 2019; de Vries et al., 2020; Gallotto et al., 2020; Klimesch, 2012; Jensen & Mazaheri, 2010). From this perspective, a decrease in alpha power likely facilitated the extraction of flexible word order rules by suppressing task-irrelevant input and optimising cortical communication in a selectively precise manner, promoting the encoding and consolidation of non-canonical grammatical rules. This interpretation is also supported by the observation that alpha power was lower for flexible relative to fixed word order rules, particularly at the beginning of the learning task (Figure 5D).

We also found that an increase in theta power predicted performance for flexible but not fixed word orders; however, theta did not interact with the aperiodic exponent to predict behavioural performance. Theta oscillations have been proposed to combine linguistic input into successively more complex representations, establishing relations between (non-adjacent) elements in a sentence (Covington & Duff, 2016; Cross et al., 2018). The positive association between theta power and performance for flexible word orders may reflect the learning and integration of non-adjacent rules, which involves the decoding and combination of words that are non-adjacent in a sentence. Indeed, such theta effects have been reported during native sentence processing (Lam et al., 2016). These effects are also consistent with the general memory literature: retrieval of language (e.g., single words), and shape/face stimuli elicit higher theta synchronisation (Bastiaansen et al., 2002; Klimesch et al., 2008; Klimesch et al., 2010; Mormann et al., 2005; Osipova et al., 2006), with these effects manifesting over medial temporal and prefrontal cortices (Guderian & Düzel, 2005), indexing the activation of relevant memory traces and executive control processes, respectively.

### Functional relevance of aperiodic activity in language and higher-order cognition

Our analysis revealed a link between aperiodic activity during language learning and performance on a grammaticality judgement task. This finding is consistent with previous studies demonstrating the influence of aperiodic activity on a range of cognitive computations, including processing speed (Ouyang et al., 2020), memory (Sheehan et al., 2018) and prediction in language (Dave et al., 2018). From a neurophysiological perspective, 1/*f*-like neural activity has been proposed to encode information relating to intrinsic brain function (Muthukumaraswamy & Liley, 2018), including the balance between excitation/inhibition (Gao et al., 2017), likely reflecting glutamate and GABA synaptic inputs into inter- and intra-cortical networks (Dave et al., 2018; Gao et al., 2017). Based on this perspective, Dave et al. (2018) argued that aperiodic activity influences prediction in language by modulating the strength of predictions of upcoming linguistic information via population spiking synchrony (Engel et al., 2001). This interpretation applies to our finding that aperiodic and beta activity showed a negative association with performance for fixed and flexible word orders: an increase in beta power predicted more sensitive behavioural responses for fixed sentences, while reduced beta predicted performance for flexible word orders. These findings can be explained by integrating two perspectives: the “spectral fingerprints” hypothesis (Hanslmayr & Staudigl, 2014; Keitel & Gross, 2016; Siegel et al., 2012; Watrous et al., 2015; Womelsdorf et al., 2014) and generalised predictive coding (Friston, 2010, 2018, 2019).

The “spectral fingerprints” hypothesis argues that power changes in different frequency bands reflect distinct stages of memory and information processing (Fellner et al., 2019; Keitel & Gross, 2016), rather than reflecting a “spectral tilt” between lower and higher frequencies. For example, decreases in alpha/beta and increases in gamma power during memory retrieval occur on different temporal scales and in different brain areas, providing evidence against proposals that a change in the tilt of the power spectrum solely drives memory computations (Fellner et al., 2019). Further, increases in high frequency gamma activity have been proposed to reflect the propagation of bottom-up sensory signals (Lewis et al., 2015; Richter, Thompson, Bosman, & Fries, 2017), while a decrease in alpha/beta power is thought to index prediction errors (Bressler & Richter, 2015; Friston, 2019; Samaha, Bauer, Cimaroli, & Postle, 2015). From this perspective, a steeper 1/*f* slope may reflect the maintenance of top-down predictions that allow comprehenders to generate expectations for incoming stimuli, thus minimizing prediction error at lower levels of the cortical hierarchy. This interpretation also holds for interactions observed with aperiodic and oscillatory activity in the alpha and beta bands, and as such, provides evidence that 1/*f*-like activity may partially reflect cortical excitability across the frequency spectrum that serves to minimize prediction error during language learning and sentence processing.

### Conclusions and Future Directions

Taken together, we have demonstrated that oscillatory and aperiodic activity jointly predict the learning of higher-order language. There are, of course, several open questions that arise from these results. For example, how do interactions between oscillatory and aperiodic activity relate to individual differences in atypical populations, such as those with schizophrenia and age-related pathologies, including Alzheimer’s disease? Previous research has shown that cognitive deficits characteristic of schizophrenia may be better explained by changes in the 1/*f* slope than irregularities in the canonical frequency bands (Peterson et al., 2018), and that 1/*f* activity mediates age-related deficits in working memory (Voytek et al., 2015); however, the interaction between aperiodic and oscillatory activity during more complex cognitive computations, such as sequence learning and language processing, remains less well known. While we attempt to address the relationship between aperiodic and oscillatory activity during higher-order language learning, future work would benefit from examining how and if these interactions emerge in (age-related) pathologies, and whether patterns of aperiodic and oscillatory activity during language learning and sentence processing are generated by specific neuroanatomical networks. Such work will provide a better understanding of the neurobiology of cognition in both health and disease.

## Acknowledgements

Preparation of this work was supported by Australian Commonwealth Government funding awarded to ZRC and AWC under the Research Training Program. IB-S is supported by an Australian Research Council Future Fellowship (FT160100437). AWC is supported by the Three Springs Foundation. We thank Isabella Sharrad, Lena Zou-Williams, Erica Wilkinson, Nicole Vass and Angela Osborn for help with data collection. Thank you also to the participants.

## Notes

**Conflict of interest statement:** The authors declare no competing financial interests.

### Competing Interest Statement

The authors have declared no competing interest.

