## Supplementary Material for "Oscillatory and aperiodic neural activity jointly predict language learning"

Table S1: Parameters from model produced by the call `lmer(formula = dprime ~ delta_dif_log * dif_bexp * type * sag * lat + (1 | subj), data = filter(dprime_summary_heh, method == "irasa"), REML = TRUE, control = lmerControl(optimizer = "bobyqa", calc.derivs = TRUE))`

| | Estimate | Std. Error | df | t value | Pr(>\|t\|) | sd subj |
|---:|---:|---:|---:|---:|---:|---:|
| (Intercept) | 1 | 0.17 | 34 | 6.1 | 7.3e-07 | 0.98 |
| delta_dif_log | -0.079 | 0.22 | 5.3e+02 | -0.36 | 0.72 | |
| dif_bexp | 0.61 | 0.5 | 5.3e+02 | 1.2 | 0.22 | |
| type[fixed] | 0.041 | 0.027 | 5e+02 | 1.5 | 0.12 | |
| sag[anterior] | -0.012 | 0.04 | 5e+02 | -0.31 | 0.75 | |
| sag[central] | 0.00097 | 0.038 | 5e+02 | 0.025 | 0.98 | |
| lat[Left] | -0.011 | 0.037 | 5.1e+02 | -0.3 | 0.76 | |
| lat[midline] | 0.0086 | 0.045 | 5.1e+02 | 0.19 | 0.85 | |
| delta_dif_log:dif_bexp | 1.9 | 1.8 | 5.2e+02 | 1 | 0.29 | |
| delta_dif_log:type[fixed] | 0.51 | 0.11 | 5e+02 | 4.7 | 3.5e-06 | |
| dif_bexp:type[fixed] | -1.4 | 0.29 | 5e+02 | -4.7 | 3.4e-06 | |
| delta_dif_log:sag[anterior] | 0.0079 | 0.15 | 5e+02 | 0.054 | 0.96 | |
| delta_dif_log:sag[central] | -0.0078 | 0.15 | 5e+02 | -0.052 | 0.96 | |
| dif_bexp:sag[anterior] | 0.1 | 0.49 | 5e+02 | 0.21 | 0.83 | |
| dif_bexp:sag[central] | 0.23 | 0.41 | 5e+02 | 0.56 | 0.58 | |
| type[fixed]:sag[anterior] | -0.029 | 0.038 | 5e+02 | -0.76 | 0.45 | |
| type[fixed]:sag[central] | 0.02 | 0.038 | 5e+02 | 0.52 | 0.6 | |
| delta_dif_log:lat[Left] | -0.065 | 0.15 | 5e+02 | -0.43 | 0.67 | |
| delta_dif_log:lat[midline] | 0.0091 | 0.15 | 5e+02 | 0.06 | 0.95 | |
| dif_bexp:lat[Left] | 0.23 | 0.42 | 5e+02 | 0.54 | 0.59 | |
| dif_bexp:lat[midline] | -0.27 | 0.47 | 5e+02 | -0.59 | 0.56 | |
| type[fixed]:lat[Left] | 0.011 | 0.035 | 5e+02 | 0.31 | 0.76 | |
| type[fixed]:lat[midline] | -0.028 | 0.041 | 5e+02 | -0.68 | 0.5 | |
| sag[anterior]:lat[Left] | -0.0033 | 0.05 | 5e+02 | -0.065 | 0.95 | |
| sag[central]:lat[Left] | -0.0017 | 0.05 | 5e+02 | -0.034 | 0.97 | |
| sag[anterior]:lat[midline] | -0.0034 | 0.06 | 5e+02 | -0.056 | 0.96 | |
| sag[central]:lat[midline] | 0.008 | 0.062 | 5e+02 | 0.13 | 0.9 | |
| delta_dif_log:dif_bexp:type[fixed] | 4.8 | 1.2 | 5e+02 | 3.9 | 0.00011 | |
| delta_dif_log:dif_bexp:sag[anterior] | 3.5 | 2 | 5e+02 | 1.8 | 0.081 | |
| delta_dif_log:dif_bexp:sag[central] | -1.2 | 1.6 | 5e+02 | -0.74 | 0.46 | |
| delta_dif_log:type[fixed]:sag[anterior] | 0.13 | 0.15 | 5e+02 | 0.9 | 0.37 | |
| delta_dif_log:type[fixed]:sag[central] | 0.049 | 0.15 | 5e+02 | 0.33 | 0.74 | |
| dif_bexp:type[fixed]:sag[anterior] | -0.32 | 0.46 | 5e+02 | -0.69 | 0.49 | |
| dif_bexp:type[fixed]:sag[central] | 0.16 | 0.41 | 5e+02 | 0.38 | 0.7 | |
| delta_dif_log:dif_bexp:lat[Left] | 2.7 | 1.9 | 5e+02 | 1.4 | 0.16 | |
| delta_dif_log:dif_bexp:lat[midline] | -1.1 | 1.7 | 5e+02 | -0.66 | 0.51 | |
| delta_dif_log:type[fixed]:lat[Left] | 0.19 | 0.15 | 5e+02 | 1.3 | 0.2 | |
| delta_dif_log:type[fixed]:lat[midline] | -0.21 | 0.15 | 5e+02 | -1.4 | 0.16 | |
| dif_bexp:type[fixed]:lat[Left] | 0.51 | 0.41 | 5e+02 | 1.2 | 0.21 | |
| dif_bexp:type[fixed]:lat[midline] | -0.31 | 0.44 | 5e+02 | -0.71 | 0.48 | |
| delta_dif_log:sag[anterior]:lat[Left] | 0.048 | 0.21 | 5e+02 | 0.23 | 0.82 | |
| delta_dif_log:sag[central]:lat[Left] | -0.036 | 0.21 | 5e+02 | -0.17 | 0.87 | |
| delta_dif_log:sag[anterior]:lat[midline] | -0.022 | 0.21 | 5e+02 | -0.1 | 0.92 | |
| delta_dif_log:sag[central]:lat[midline] | 0.036 | 0.21 | 5e+02 | 0.17 | 0.87 | |
| dif_bexp:sag[anterior]:lat[Left] | 0.17 | 0.64 | 5e+02 | 0.27 | 0.79 | |
| dif_bexp:sag[central]:lat[Left] | 0.00089 | 0.59 | 5e+02 | 0.0015 | 1 | |
| dif_bexp:sag[anterior]:lat[midline] | -0.29 | 0.73 | 5e+02 | -0.39 | 0.69 | |
| dif_bexp:sag[central]:lat[midline] | 0.099 | 0.62 | 5e+02 | 0.16 | 0.87 | |
| type[fixed]:sag[anterior]:lat[Left] | -0.0056 | 0.05 | 5e+02 | -0.11 | 0.91 | |
| type[fixed]:sag[central]:lat[Left] | 0.0067 | 0.05 | 5e+02 | 0.13 | 0.89 | |
| type[fixed]:sag[anterior]:lat[midline] | 0.0094 | 0.059 | 5e+02 | 0.16 | 0.87 | |
| type[fixed]:sag[central]:lat[midline] | -0.012 | 0.061 | 5e+02 | -0.19 | 0.85 | |
| delta_dif_log:dif_bexp:type[fixed]:sag[anterior] | 2.2 | 1.9 | 5e+02 | 1.2 | 0.25 | |
| delta_dif_log:dif_bexp:type[fixed]:sag[central] | 0.51 | 1.6 | 5e+02 | 0.32 | 0.75 | |
| delta_dif_log:dif_bexp:type[fixed]:lat[Left] | 2 | 1.9 | 5e+02 | 1.1 | 0.28 | |
| delta_dif_log:dif_bexp:type[fixed]:lat[midline] | -2.6 | 1.7 | 5e+02 | -1.6 | 0.11 | |
| delta_dif_log:dif_bexp:sag[anterior]:lat[Left] | 3 | 2.9 | 5e+02 | 1 | 0.3 | |
| delta_dif_log:dif_bexp:sag[central]:lat[Left] | -0.77 | 2.4 | 5e+02 | -0.32 | 0.75 | |
| delta_dif_log:dif_bexp:sag[anterior]:lat[midline] | -0.71 | 2.7 | 5e+02 | -0.26 | 0.79 | |
| delta_dif_log:dif_bexp:sag[central]:lat[midline] | -0.34 | 2.2 | 5e+02 | -0.15 | 0.88 | |
| delta_dif_log:type[fixed]:sag[anterior]:lat[Left] | -0.1 | 0.2 | 5e+02 | -0.5 | 0.62 | |
| delta_dif_log:type[fixed]:sag[central]:lat[Left] | 0.11 | 0.21 | 5e+02 | 0.54 | 0.59 | |
| delta_dif_log:type[fixed]:sag[anterior]:lat[midline] | 0.089 | 0.21 | 5e+02 | 0.43 | 0.67 | |
| delta_dif_log:type[fixed]:sag[central]:lat[midline] | -0.094 | 0.21 | 5e+02 | -0.44 | 0.66 | |
| dif_bexp:type[fixed]:sag[anterior]:lat[Left] | 0.34 | 0.63 | 5e+02 | 0.54 | 0.59 | |
| dif_bexp:type[fixed]:sag[central]:lat[Left] | 0.19 | 0.58 | 5e+02 | 0.32 | 0.75 | |
| dif_bexp:type[fixed]:sag[anterior]:lat[midline] | -0.24 | 0.72 | 5e+02 | -0.33 | 0.74 | |
| dif_bexp:type[fixed]:sag[central]:lat[midline] | -0.15 | 0.61 | 5e+02 | -0.24 | 0.81 | |
| delta_dif_log:dif_bexp:type[fixed]:sag[anterior]:lat[Left] | 2.5 | 2.9 | 5e+02 | 0.85 | 0.4 | |
| delta_dif_log:dif_bexp:type[fixed]:sag[central]:lat[Left] | -0.94 | 2.4 | 5e+02 | -0.39 | 0.7 | |
| delta_dif_log:dif_bexp:type[fixed]:sag[anterior]:lat[midline] | -0.25 | 2.7 | 5e+02 | -0.092 | 0.93 | |
| delta_dif_log:dif_bexp:type[fixed]:sag[central]:lat[midline] | -0.037 | 2.2 | 5e+02 | -0.017 | 0.99 | |

Table S2: Parameters from model produced by the call `lmer(formula = dprime ~ theta_dif_log * dif_bexp * type * sag * lat + (1 | subj), data = filter(dprime_summary_heh, method == "irasa"), REML = TRUE, control = lmerControl(optimizer = "bobyqa", calc.derivs = TRUE))`

| | Estimate | Std. Error | df | t value | Pr(>\|t\|) | sd subj |
|---:|---:|---:|---:|---:|---:|---:|
| (Intercept) | 1 | 0.17 | 37 | 5.9 | 8.9e-07 | 0.98 |
| theta_dif_log | -0.035 | 0.21 | 5.3e+02 | -0.17 | 0.87 | |
| dif_bexp | 0.37 | 0.57 | 5.3e+02 | 0.65 | 0.52 | |
| type[fixed] | 0.13 | 0.032 | 5e+02 | 4.2 | 3.7e-05 | |
| sag[anterior] | -0.0076 | 0.049 | 5.2e+02 | -0.15 | 0.88 | |
| sag[central] | 0.0013 | 0.044 | 5e+02 | 0.03 | 0.98 | |
| lat[Left] | 0.011 | 0.048 | 5.1e+02 | 0.22 | 0.82 | |
| lat[midline] | -0.002 | 0.051 | 5.2e+02 | -0.04 | 0.97 | |
| theta_dif_log:dif_bexp | -0.46 | 1.6 | 5.3e+02 | -0.29 | 0.78 | |
| theta_dif_log:type[fixed] | 0.21 | 0.092 | 5e+02 | 2.3 | 0.021 | |
| dif_bexp:type[fixed] | -0.68 | 0.33 | 5e+02 | -2.1 | 0.041 | |
| theta_dif_log:sag[anterior] | -0.055 | 0.13 | 5e+02 | -0.41 | 0.68 | |
| theta_dif_log:sag[central] | 0.015 | 0.13 | 5e+02 | 0.12 | 0.91 | |
| dif_bexp:sag[anterior] | 0.17 | 0.49 | 5e+02 | 0.35 | 0.73 | |
| dif_bexp:sag[central] | 0.24 | 0.46 | 5e+02 | 0.52 | 0.6 | |
| type[fixed]:sag[anterior] | 0.0052 | 0.041 | 5e+02 | 0.13 | 0.9 | |
| type[fixed]:sag[central] | 0.014 | 0.043 | 5e+02 | 0.32 | 0.75 | |
| theta_dif_log:lat[Left] | -0.0029 | 0.13 | 5e+02 | -0.023 | 0.98 | |
| theta_dif_log:lat[midline] | 4.5e-05 | 0.13 | 5e+02 | 0.00035 | 1 | |
| dif_bexp:lat[Left] | 0.027 | 0.53 | 5e+02 | 0.05 | 0.96 | |
| dif_bexp:lat[midline] | -0.1 | 0.47 | 5.1e+02 | -0.21 | 0.83 | |
| type[fixed]:lat[Left] | 0.026 | 0.046 | 5e+02 | 0.57 | 0.57 | |
| type[fixed]:lat[midline] | -0.043 | 0.044 | 5e+02 | -0.97 | 0.33 | |
| sag[anterior]:lat[Left] | -0.0072 | 0.059 | 5e+02 | -0.12 | 0.9 | |
| sag[central]:lat[Left] | 0.007 | 0.064 | 5e+02 | 0.11 | 0.91 | |
| sag[anterior]:lat[midline] | 0.0037 | 0.059 | 5e+02 | 0.063 | 0.95 | |
| sag[central]:lat[midline] | 0.012 | 0.059 | 5e+02 | 0.21 | 0.83 | |
| theta_dif_log:dif_bexp:type[fixed] | 0.59 | 0.98 | 5e+02 | 0.61 | 0.55 | |
| theta_dif_log:dif_bexp:sag[anterior] | -0.38 | 1.6 | 5e+02 | -0.23 | 0.82 | |
| theta_dif_log:dif_bexp:sag[central] | 0.51 | 1.3 | 5e+02 | 0.4 | 0.69 | |
| theta_dif_log:type[fixed]:sag[anterior] | 0.07 | 0.13 | 5e+02 | 0.53 | 0.59 | |
| theta_dif_log:type[fixed]:sag[central] | -0.0079 | 0.13 | 5e+02 | -0.062 | 0.95 | |
| dif_bexp:type[fixed]:sag[anterior] | -0.13 | 0.48 | 5e+02 | -0.27 | 0.79 | |
| dif_bexp:type[fixed]:sag[central] | -0.099 | 0.46 | 5e+02 | -0.22 | 0.83 | |
| theta_dif_log:dif_bexp:lat[Left] | -0.67 | 1.5 | 5e+02 | -0.46 | 0.65 | |
| theta_dif_log:dif_bexp:lat[midline] | 0.6 | 1.3 | 5e+02 | 0.44 | 0.66 | |
| theta_dif_log:type[fixed]:lat[Left] | 0.035 | 0.13 | 5e+02 | 0.27 | 0.78 | |
| theta_dif_log:type[fixed]:lat[midline] | -0.036 | 0.13 | 5e+02 | -0.28 | 0.78 | |
| dif_bexp:type[fixed]:lat[Left] | 0.13 | 0.51 | 5e+02 | 0.25 | 0.8 | |
| dif_bexp:type[fixed]:lat[midline] | -0.02 | 0.43 | 5e+02 | -0.046 | 0.96 | |
| theta_dif_log:sag[anterior]:lat[Left] | -0.094 | 0.18 | 5e+02 | -0.51 | 0.61 | |
| theta_dif_log:sag[central]:lat[Left] | 0.061 | 0.18 | 5e+02 | 0.34 | 0.73 | |
| theta_dif_log:sag[anterior]:lat[midline] | 0.051 | 0.19 | 5e+02 | 0.27 | 0.79 | |
| theta_dif_log:sag[central]:lat[midline] | 0.0055 | 0.18 | 5e+02 | 0.031 | 0.98 | |
| dif_bexp:sag[anterior]:lat[Left] | -0.17 | 0.75 | 5e+02 | -0.23 | 0.82 | |
| dif_bexp:sag[central]:lat[Left] | 0.015 | 0.72 | 5e+02 | 0.021 | 0.98 | |
| dif_bexp:sag[anterior]:lat[midline] | 0.063 | 0.65 | 5e+02 | 0.097 | 0.92 | |
| dif_bexp:sag[central]:lat[midline] | -0.003 | 0.59 | 5e+02 | -0.0051 | 1 | |
| type[fixed]:sag[anterior]:lat[Left] | -0.0016 | 0.059 | 5e+02 | -0.027 | 0.98 | |
| type[fixed]:sag[central]:lat[Left] | -0.003 | 0.064 | 5e+02 | -0.047 | 0.96 | |
| type[fixed]:sag[anterior]:lat[midline] | 0.00046 | 0.057 | 5e+02 | 0.008 | 0.99 | |
| type[fixed]:sag[central]:lat[midline] | -0.0094 | 0.058 | 5e+02 | -0.16 | 0.87 | |
| theta_dif_log:dif_bexp:type[fixed]:sag[anterior] | -2 | 1.6 | 5e+02 | -1.2 | 0.22 | |
| theta_dif_log:dif_bexp:type[fixed]:sag[central] | 0.39 | 1.3 | 5e+02 | 0.31 | 0.76 | |
| theta_dif_log:dif_bexp:type[fixed]:lat[Left] | -1.1 | 1.4 | 5e+02 | -0.79 | 0.43 | |
| theta_dif_log:dif_bexp:type[fixed]:lat[midline] | -0.2 | 1.3 | 5e+02 | -0.15 | 0.88 | |
| theta_dif_log:dif_bexp:sag[anterior]:lat[Left] | -2.3 | 2.4 | 5e+02 | -0.94 | 0.35 | |
| theta_dif_log:dif_bexp:sag[central]:lat[Left] | 0.98 | 1.9 | 5e+02 | 0.52 | 0.61 | |
| theta_dif_log:dif_bexp:sag[anterior]:lat[midline] | 1.9 | 2.2 | 5e+02 | 0.88 | 0.38 | |
| theta_dif_log:dif_bexp:sag[central]:lat[midline] | -1.2 | 1.8 | 5e+02 | -0.66 | 0.51 | |
| theta_dif_log:type[fixed]:sag[anterior]:lat[Left] | -0.047 | 0.18 | 5e+02 | -0.26 | 0.8 | |
| theta_dif_log:type[fixed]:sag[central]:lat[Left] | 0.0027 | 0.18 | 5e+02 | 0.015 | 0.99 | |
| theta_dif_log:type[fixed]:sag[anterior]:lat[midline] | -0.024 | 0.19 | 5e+02 | -0.13 | 0.9 | |
| theta_dif_log:type[fixed]:sag[central]:lat[midline] | -0.0091 | 0.18 | 5e+02 | -0.051 | 0.96 | |
| dif_bexp:type[fixed]:sag[anterior]:lat[Left] | -0.025 | 0.74 | 5e+02 | -0.034 | 0.97 | |
| dif_bexp:type[fixed]:sag[central]:lat[Left] | 0.31 | 0.72 | 5e+02 | 0.43 | 0.67 | |
| dif_bexp:type[fixed]:sag[anterior]:lat[midline] | 0.15 | 0.64 | 5e+02 | 0.23 | 0.82 | |
| dif_bexp:type[fixed]:sag[central]:lat[midline] | -0.24 | 0.58 | 5e+02 | -0.41 | 0.68 | |
| theta_dif_log:dif_bexp:type[fixed]:sag[anterior]:lat[Left] | -1.2 | 2.4 | 5e+02 | -0.51 | 0.61 | |
| theta_dif_log:dif_bexp:type[fixed]:sag[central]:lat[Left] | 0.71 | 1.9 | 5e+02 | 0.38 | 0.71 | |
| theta_dif_log:dif_bexp:type[fixed]:sag[anterior]:lat[midline] | -0.21 | 2.2 | 5e+02 | -0.098 | 0.92 | |
| theta_dif_log:dif_bexp:type[fixed]:sag[central]:lat[midline] | 0.3 | 1.8 | 5e+02 | 0.17 | 0.87 | |

Table S3: Parameters from model produced by the call `lmer(formula = dprime ~ alpha_dif_log * dif_bexp * type * sag * lat + (1 | subj), data = filter(dprime_summary_heh, method == "irasa"), REML = TRUE, control = lmerControl(optimizer = "bobyqa", calc.derivs = TRUE))`

| | Estimate | Std. Error | df | t value | Pr(>\|t\|) | sd subj |
|---:|---:|---:|---:|---:|---:|---:|
| (Intercept) | 0.97 | 0.17 | 35 | 5.8 | 1.2e-06 | 0.94 |
| alpha_dif_log | -0.016 | 0.15 | 5.2e+02 | -0.1 | 0.92 | |
| dif_bexp | 0.21 | 0.52 | 5.3e+02 | 0.41 | 0.68 | |
| type[fixed] | 0.16 | 0.028 | 5e+02 | 5.6 | 4e-08 | |
| sag[anterior] | 0.0099 | 0.041 | 5.1e+02 | 0.24 | 0.81 | |
| sag[central] | -0.011 | 0.039 | 5e+02 | -0.29 | 0.77 | |
| lat[Left] | -0.0017 | 0.04 | 5e+02 | -0.043 | 0.97 | |
| lat[midline] | -0.0037 | 0.041 | 5.1e+02 | -0.09 | 0.93 | |
| alpha_dif_log:dif_bexp | -3.3 | 0.9 | 5.2e+02 | -3.6 | 0.00029 | |
| alpha_dif_log:type[fixed] | 0.069 | 0.063 | 5e+02 | 1.1 | 0.28 | |
| dif_bexp:type[fixed] | -0.7 | 0.31 | 5e+02 | -2.2 | 0.025 | |
| alpha_dif_log:sag[anterior] | 0.061 | 0.088 | 5e+02 | 0.7 | 0.49 | |
| alpha_dif_log:sag[central] | 0.052 | 0.09 | 5e+02 | 0.58 | 0.56 | |
| dif_bexp:sag[anterior] | 0.1 | 0.45 | 5e+02 | 0.23 | 0.82 | |
| dif_bexp:sag[central] | 0.092 | 0.44 | 5e+02 | 0.21 | 0.83 | |
| type[fixed]:sag[anterior] | 0.011 | 0.039 | 5e+02 | 0.27 | 0.79 | |
| type[fixed]:sag[central] | 0.021 | 0.038 | 5e+02 | 0.55 | 0.58 | |
| alpha_dif_log:lat[Left] | -0.0076 | 0.087 | 5e+02 | -0.088 | 0.93 | |
| alpha_dif_log:lat[midline] | -0.039 | 0.089 | 5e+02 | -0.44 | 0.66 | |
| dif_bexp:lat[Left] | -0.24 | 0.45 | 5e+02 | -0.52 | 0.6 | |
| dif_bexp:lat[midline] | 0.38 | 0.42 | 5e+02 | 0.89 | 0.38 | |
| type[fixed]:lat[Left] | -0.0075 | 0.039 | 5e+02 | -0.19 | 0.85 | |
| type[fixed]:lat[midline] | -0.033 | 0.038 | 5e+02 | -0.86 | 0.39 | |
| sag[anterior]:lat[Left] | 0.0096 | 0.055 | 5e+02 | 0.17 | 0.86 | |
| sag[central]:lat[Left] | 0.004 | 0.055 | 5e+02 | 0.073 | 0.94 | |
| sag[anterior]:lat[midline] | -0.0046 | 0.053 | 5e+02 | -0.087 | 0.93 | |
| sag[central]:lat[midline] | -0.0063 | 0.053 | 5e+02 | -0.12 | 0.91 | |
| alpha_dif_log:dif_bexp:type[fixed] | 2 | 0.58 | 5e+02 | 3.5 | 0.00061 | |
| alpha_dif_log:dif_bexp:sag[anterior] | -0.79 | 0.86 | 5e+02 | -0.92 | 0.36 | |
| alpha_dif_log:dif_bexp:sag[central] | -0.8 | 0.86 | 5e+02 | -0.93 | 0.35 | |
| alpha_dif_log:type[fixed]:sag[anterior] | -0.043 | 0.087 | 5e+02 | -0.49 | 0.62 | |
| alpha_dif_log:type[fixed]:sag[central] | 0.029 | 0.089 | 5e+02 | 0.32 | 0.75 | |
| dif_bexp:type[fixed]:sag[anterior] | 0.4 | 0.43 | 5e+02 | 0.92 | 0.36 | |
| dif_bexp:type[fixed]:sag[central] | 0.11 | 0.43 | 5e+02 | 0.26 | 0.79 | |
| alpha_dif_log:dif_bexp:lat[Left] | -0.86 | 0.92 | 5e+02 | -0.93 | 0.35 | |
| alpha_dif_log:dif_bexp:lat[midline] | 0.91 | 0.73 | 5e+02 | 1.3 | 0.21 | |
| alpha_dif_log:type[fixed]:lat[Left] | 0.044 | 0.087 | 5e+02 | 0.51 | 0.61 | |
| alpha_dif_log:type[fixed]:lat[midline] | -0.022 | 0.088 | 5e+02 | -0.25 | 0.8 | |
| dif_bexp:type[fixed]:lat[Left] | 0.46 | 0.44 | 5e+02 | 1 | 0.3 | |
| dif_bexp:type[fixed]:lat[midline] | -0.42 | 0.41 | 5e+02 | -1 | 0.3 | |
| alpha_dif_log:sag[anterior]:lat[Left] | -0.033 | 0.12 | 5e+02 | -0.28 | 0.78 | |
| alpha_dif_log:sag[central]:lat[Left] | -0.024 | 0.12 | 5e+02 | -0.19 | 0.85 | |
| alpha_dif_log:sag[anterior]:lat[midline] | 0.053 | 0.12 | 5e+02 | 0.44 | 0.66 | |
| alpha_dif_log:sag[central]:lat[midline] | 0.015 | 0.12 | 5e+02 | 0.12 | 0.9 | |
| dif_bexp:sag[anterior]:lat[Left] | 0.16 | 0.64 | 5e+02 | 0.25 | 0.8 | |
| dif_bexp:sag[central]:lat[Left] | -0.48 | 0.65 | 5e+02 | -0.75 | 0.46 | |
| dif_bexp:sag[anterior]:lat[midline] | 0.23 | 0.61 | 5e+02 | 0.38 | 0.7 | |
| dif_bexp:sag[central]:lat[midline] | 0.42 | 0.6 | 5e+02 | 0.7 | 0.49 | |
| type[fixed]:sag[anterior]:lat[Left] | 0.014 | 0.055 | 5e+02 | 0.25 | 0.8 | |
| type[fixed]:sag[central]:lat[Left] | -0.023 | 0.055 | 5e+02 | -0.41 | 0.68 | |
| type[fixed]:sag[anterior]:lat[midline] | 0.015 | 0.053 | 5e+02 | 0.28 | 0.78 | |
| type[fixed]:sag[central]:lat[midline] | -0.0066 | 0.053 | 5e+02 | -0.12 | 0.9 | |
| alpha_dif_log:dif_bexp:type[fixed]:sag[anterior] | 0.22 | 0.83 | 5e+02 | 0.27 | 0.79 | |
| alpha_dif_log:dif_bexp:type[fixed]:sag[central] | -0.095 | 0.84 | 5e+02 | -0.11 | 0.91 | |
| alpha_dif_log:dif_bexp:type[fixed]:lat[Left] | -1.3 | 0.89 | 5e+02 | -1.5 | 0.15 | |
| alpha_dif_log:dif_bexp:type[fixed]:lat[midline] | -0.57 | 0.71 | 5e+02 | -0.8 | 0.42 | |
| alpha_dif_log:dif_bexp:sag[anterior]:lat[Left] | 0.48 | 1.3 | 5e+02 | 0.38 | 0.71 | |
| alpha_dif_log:dif_bexp:sag[central]:lat[Left] | -0.51 | 1.4 | 5e+02 | -0.37 | 0.71 | |
| alpha_dif_log:dif_bexp:sag[anterior]:lat[midline] | 0.21 | 1.1 | 5e+02 | 0.19 | 0.85 | |
| alpha_dif_log:dif_bexp:sag[central]:lat[midline] | 0.21 | 1 | 5e+02 | 0.2 | 0.84 | |
| alpha_dif_log:type[fixed]:sag[anterior]:lat[Left] | 0.056 | 0.12 | 5e+02 | 0.47 | 0.64 | |
| alpha_dif_log:type[fixed]:sag[central]:lat[Left] | 0.027 | 0.12 | 5e+02 | 0.22 | 0.82 | |
| alpha_dif_log:type[fixed]:sag[anterior]:lat[midline] | -0.076 | 0.12 | 5e+02 | -0.63 | 0.53 | |
| alpha_dif_log:type[fixed]:sag[central]:lat[midline] | -0.034 | 0.12 | 5e+02 | -0.28 | 0.78 | |
| dif_bexp:type[fixed]:sag[anterior]:lat[Left] | 0.37 | 0.63 | 5e+02 | 0.59 | 0.56 | |
| dif_bexp:type[fixed]:sag[central]:lat[Left] | -0.12 | 0.64 | 5e+02 | -0.18 | 0.86 | |
| dif_bexp:type[fixed]:sag[anterior]:lat[midline] | -0.4 | 0.6 | 5e+02 | -0.67 | 0.5 | |
| dif_bexp:type[fixed]:sag[central]:lat[midline] | -0.16 | 0.6 | 5e+02 | -0.27 | 0.79 | |
| alpha_dif_log:dif_bexp:type[fixed]:sag[anterior]:lat[Left] | 0.35 | 1.3 | 5e+02 | 0.28 | 0.78 | |
| alpha_dif_log:dif_bexp:type[fixed]:sag[central]:lat[Left] | -1.5 | 1.4 | 5e+02 | -1.1 | 0.28 | |
| alpha_dif_log:dif_bexp:type[fixed]:sag[anterior]:lat[midline] | -0.51 | 1.1 | 5e+02 | -0.48 | 0.63 | |
| alpha_dif_log:dif_bexp:type[fixed]:sag[central]:lat[midline] | 0.088 | 1 | 5e+02 | 0.085 | 0.93 | |

Table S4: Parameters from model produced by the call `lmer(formula = dprime ~ beta_dif_log * dif_bexp * type * sag * lat + (1 | subj), data = filter(dprime_summary_heh, method == "irasa"), REML = TRUE, control = lmerControl(optimizer = "bobyqa", calc.derivs = TRUE))`

| | Estimate | Std. Error | df | t value | Pr(>\|t\|) | sd subj |
|---|---|---|---|---|---|---|
| (Intercept) | 0.74 | 0.21 | 74 | 3.5 | 0.00069 | 0.96 |
| beta_dif_log | -0.35 | 0.16 | 5.3e+02 | -2.2 | 0.029 | |
| dif_bexp | 2.1 | 1.1 | 5.2e+02 | 1.9 | 0.065 | |
| type[fixed] | 0.48 | 0.064 | 5e+02 | 7.6 | 1.5e-13 | |
| sag[anterior] | 0.027 | 0.083 | 5e+02 | 0.32 | 0.75 | |
| sag[central] | 0.022 | 0.082 | 5e+02 | 0.26 | 0.79 | |
| lat[Left] | 0.025 | 0.082 | 5e+02 | 0.31 | 0.76 | |
| lat[midline] | -0.016 | 0.09 | 5e+02 | -0.17 | 0.86 | |
| beta_dif_log:dif_bexp | 1.4 | 1.3 | 5.2e+02 | 1.1 | 0.27 | |
| beta_dif_log:type[fixed] | 0.47 | 0.072 | 5e+02 | 6.6 | 1.4e-10 | |
| dif_bexp:type[fixed] | 2.7 | 0.68 | 5e+02 | 4 | 7.4e-05 | |
| beta_dif_log:sag[anterior] | 0.027 | 0.097 | 5e+02 | 0.28 | 0.78 | |
| beta_dif_log:sag[central] | 0.021 | 0.094 | 5e+02 | 0.22 | 0.82 | |
| dif_bexp:sag[anterior] | 0.25 | 0.95 | 5e+02 | 0.26 | 0.79 | |
| dif_bexp:sag[central] | 0.31 | 0.91 | 5e+02 | 0.34 | 0.74 | |
| type[fixed]:sag[anterior] | -0.064 | 0.08 | 5e+02 | -0.81 | 0.42 | |
| type[fixed]:sag[central] | -0.048 | 0.081 | 5e+02 | -0.59 | 0.55 | |
| beta_dif_log:lat[Left] | 0.023 | 0.094 | 5e+02 | 0.25 | 0.8 | |
| beta_dif_log:lat[midline] | -0.032 | 0.1 | 5e+02 | -0.31 | 0.76 | |
| dif_bexp:lat[Left] | 0.97 | 0.97 | 5e+02 | 0.99 | 0.32 | |
| dif_bexp:lat[midline] | -0.87 | 0.91 | 5e+02 | -0.95 | 0.34 | |
| type[fixed]:lat[Left] | 0.033 | 0.082 | 5e+02 | 0.4 | 0.69 | |
| type[fixed]:lat[midline] | 0.0035 | 0.089 | 5e+02 | 0.04 | 0.97 | |
| sag[anterior]:lat[Left] | -0.025 | 0.11 | 5e+02 | -0.23 | 0.82 | |
| sag[central]:lat[Left] | -0.0048 | 0.11 | 5e+02 | -0.043 | 0.97 | |
| sag[anterior]:lat[midline] | -0.082 | 0.12 | 5e+02 | -0.67 | 0.5 | |
| sag[central]:lat[midline] | 0.015 | 0.12 | 5e+02 | 0.13 | 0.9 | |
| beta_dif_log:dif_bexp:type[fixed] | 4.3 | 0.76 | 5e+02 | 5.7 | 1.8e-08 | |
| beta_dif_log:dif_bexp:sag[anterior] | 0.11 | 1.1 | 5e+02 | 0.099 | 0.92 | |
| beta_dif_log:dif_bexp:sag[central] | 0.15 | 1 | 5e+02 | 0.15 | 0.88 | |
| beta_dif_log:type[fixed]:sag[anterior] | -0.074 | 0.094 | 5e+02 | -0.79 | 0.43 | |
| beta_dif_log:type[fixed]:sag[central] | -0.065 | 0.092 | 5e+02 | -0.7 | 0.48 | |
| dif_bexp:type[fixed]:sag[anterior] | 0.5 | 0.94 | 5e+02 | 0.53 | 0.6 | |
| dif_bexp:type[fixed]:sag[central] | 0.42 | 0.91 | 5e+02 | 0.46 | 0.65 | |
| beta_dif_log:dif_bexp:lat[Left] | 0.88 | 1.1 | 5e+02 | 0.82 | 0.41 | |
| beta_dif_log:dif_bexp:lat[midline] | -0.76 | 1 | 5e+02 | -0.73 | 0.47 | |
| beta_dif_log:type[fixed]:lat[Left] | 0.034 | 0.093 | 5e+02 | 0.37 | 0.71 | |
| beta_dif_log:type[fixed]:lat[midline] | 0.021 | 0.1 | 5e+02 | 0.21 | 0.84 | |
| dif_bexp:type[fixed]:lat[Left] | 1.4 | 0.97 | 5e+02 | 1.4 | 0.15 | |
| dif_bexp:type[fixed]:lat[midline] | -0.33 | 0.91 | 5e+02 | -0.37 | 0.71 | |
| beta_dif_log:sag[anterior]:lat[Left] | -0.037 | 0.13 | 5e+02 | -0.29 | 0.77 | |
| beta_dif_log:sag[central]:lat[Left] | -0.0057 | 0.13 | 5e+02 | -0.044 | 0.96 | |
| beta_dif_log:sag[anterior]:lat[midline] | -0.072 | 0.14 | 5e+02 | -0.5 | 0.62 | |
| beta_dif_log:sag[central]:lat[midline] | 0.018 | 0.14 | 5e+02 | 0.13 | 0.9 | |
| dif_bexp:sag[anterior]:lat[Left] | 0.35 | 1.4 | 5e+02 | 0.25 | 0.8 | |
| dif_bexp:sag[central]:lat[Left] | 0.28 | 1.4 | 5e+02 | 0.2 | 0.84 | |
| dif_bexp:sag[anterior]:lat[midline] | -0.04 | 1.3 | 5e+02 | -0.03 | 0.98 | |
| dif_bexp:sag[central]:lat[midline] | 0.14 | 1.3 | 5e+02 | 0.11 | 0.91 | |
| type[fixed]:sag[anterior]:lat[Left] | -0.017 | 0.11 | 5e+02 | -0.16 | 0.88 | |
| type[fixed]:sag[central]:lat[Left] | -0.05 | 0.11 | 5e+02 | -0.44 | 0.66 | |
| type[fixed]:sag[anterior]:lat[midline] | -0.048 | 0.12 | 5e+02 | -0.4 | 0.69 | |
| type[fixed]:sag[central]:lat[midline] | 0.058 | 0.12 | 5e+02 | 0.48 | 0.63 | |
| beta_dif_log:dif_bexp:type[fixed]:sag[anterior] | 0.68 | 1.1 | 5e+02 | 0.61 | 0.54 | |
| beta_dif_log:dif_bexp:type[fixed]:sag[central] | 0.59 | 1 | 5e+02 | 0.58 | 0.56 | |
| beta_dif_log:dif_bexp:type[fixed]:lat[Left] | 0.93 | 1.1 | 5e+02 | 0.86 | 0.39 | |
| beta_dif_log:dif_bexp:type[fixed]:lat[midline] | -0.18 | 1 | 5e+02 | -0.18 | 0.86 | |
| beta_dif_log:dif_bexp:sag[anterior]:lat[Left] | 0.35 | 1.6 | 5e+02 | 0.22 | 0.83 | |
| beta_dif_log:dif_bexp:sag[central]:lat[Left] | 0.2 | 1.6 | 5e+02 | 0.13 | 0.9 | |
| beta_dif_log:dif_bexp:sag[anterior]:lat[midline] | -0.23 | 1.6 | 5e+02 | -0.15 | 0.88 | |
| beta_dif_log:dif_bexp:sag[central]:lat[midline] | 0.35 | 1.4 | 5e+02 | 0.25 | 0.81 | |
| beta_dif_log:type[fixed]:sag[anterior]:lat[Left] | 0.0066 | 0.13 | 5e+02 | 0.052 | 0.96 | |
| beta_dif_log:type[fixed]:sag[central]:lat[Left] | -0.044 | 0.13 | 5e+02 | -0.35 | 0.73 | |
| beta_dif_log:type[fixed]:sag[anterior]:lat[midline] | -0.11 | 0.14 | 5e+02 | -0.8 | 0.42 | |
| beta_dif_log:type[fixed]:sag[central]:lat[midline] | 0.056 | 0.14 | 5e+02 | 0.41 | 0.68 | |
| dif_bexp:type[fixed]:sag[anterior]:lat[Left] | -0.14 | 1.4 | 5e+02 | -0.1 | 0.92 | |
| dif_bexp:type[fixed]:sag[central]:lat[Left] | 0.71 | 1.4 | 5e+02 | 0.51 | 0.61 | |
| dif_bexp:type[fixed]:sag[anterior]:lat[midline] | 0.39 | 1.3 | 5e+02 | 0.29 | 0.77 | |
| dif_bexp:type[fixed]:sag[central]:lat[midline] | -0.013 | 1.3 | 5e+02 | -0.01 | 0.99 | |
| beta_dif_log:dif_bexp:type[fixed]:sag[anterior]:lat[Left] | -0.66 | 1.6 | 5e+02 | -0.42 | 0.68 | |
| beta_dif_log:dif_bexp:type[fixed]:sag[central]:lat[Left] | 0.88 | 1.6 | 5e+02 | 0.56 | 0.58 | |
| beta_dif_log:dif_bexp:type[fixed]:sag[anterior]:lat[midline] | 0.65 | 1.6 | 5e+02 | 0.41 | 0.68 | |
| beta_dif_log:dif_bexp:type[fixed]:sag[central]:lat[midline] | -0.3 | 1.4 | 5e+02 | -0.21 | 0.83 | |

### Generalised Additive Mixed Model Summaries

Cross, Corcoran et al., 2022

April 2022

#### 1 A1. Aperiodic Exponent (IRASA)

| A. parametric coefficients | Estimate | Std. Error | t-value | p-value |
|---|---|---|---|---|
| (Intercept) | 0.6062 | 0.0336 | 18.0241 | $< 0.0001$ |
| bln | 0.3724 | 0.0032 | 117.7504 | $< 0.0001$ |
| Typeflexible | 0.0204 | 0.0084 | 2.4157 | 0.0157 |
| B. smooth terms | edf | Ref.df | F-value | p-value |
| ti(Trial.s) | 3.0977 | 4.0000 | 9.8477 | 0.0013 |
| ti(Lat.XY) | 2.9468 | 4.0000 | 86.2180 | $< 0.0001$ |
| ti(Sag.XY) | 3.6359 | 4.0000 | 767.8654 | $< 0.0001$ |
| ti(Trial.s):Typeflexible | 3.8706 | 4.0000 | 29.6675 | $< 0.0001$ |
| ti(Lat.XY):Typeflexible | 0.6541 | 4.0000 | 0.4726 | 0.0658 |
| ti(Sag.XY):Typeflexible | 0.9060 | 4.0000 | 2.4059 | 0.0006 |
| ti(Trial.s,Lat.XY) | 3.3853 | 16.0000 | 1.4003 | $< 0.0001$ |
| ti(Trial.s,Sag.XY) | 1.9840 | 16.0000 | 0.3256 | 0.0388 |
| ti(Lat.XY,Sag.XY) | 15.4955 | 17.0000 | 12.4173 | $< 0.0001$ |
| ti(Trial.s,Lat.XY):Typeflexible | 6.2610 | 16.0000 | 1.6455 | $< 0.0001$ |
| ti(Trial.s,Sag.XY):Typeflexible | 1.7838 | 13.0000 | 1.1318 | 0.0001 |
| ti(Lat.XY,Sag.XY):Typeflexible | 0.3968 | 17.0000 | 0.0256 | 0.2396 |
| ti(Trial.s,Lat.XY,Sag.XY) | 1.4277 | 68.0000 | 0.0349 | 0.1038 |
| ti(Trial.s,Lat.XY,Sag.XY):Typeflexible | 0.0001 | 68.0000 | 0.0000 | 0.6159 |
| s(tpnt.s,Subject) | 299.7417 | 314.0000 | 1000.9491 | $< 0.0001$ |
| s(Type,Subject) | 31.7868 | 67.0000 | 8.9784 | $< 0.0001$ |

### 2 A2. Delta Power (IRASA)

| A. parametric coefficients | Estimate | Std. Error | t-value | p-value |
|---|---|---|---|---|
| (Intercept) | 0.2364 | 0.0341 | 6.9265 | < 0.0001 |
| bln | 0.0601 | 0.0030 | 20.1406 | < 0.0001 |
| Typeflexible | 0.0299 | 0.0156 | 1.9189 | 0.0550 |
| B. smooth terms | edf | Ref.df | F-value | p-value |
| ti(Trial.s) | 3.7187 | 4.0000 | 15.9602 | < 0.0001 |
| ti(Lat.XY) | 3.6464 | 4.0000 | 144.0866 | < 0.0001 |
| ti(Sag.XY) | 3.9203 | 4.0000 | 719.9381 | < 0.0001 |
| ti(Trial.s):Typeflexible | 2.7429 | 4.0000 | 9.6596 | < 0.0001 |
| ti(Lat.XY):Typeflexible | 0.0003 | 4.0000 | 0.0000 | 0.8428 |
| ti(Sag.XY):Typeflexible | 0.9220 | 4.0000 | 2.9405 | 0.0004 |
| ti(Trial.s,Lat.XY) | 4.2900 | 16.0000 | 0.6699 | 0.0041 |
| ti(Trial.s,Sag.XY) | 5.4273 | 16.0000 | 0.9238 | 0.0030 |
| ti(Lat.XY,Sag.XY) | 16.7539 | 17.0000 | 47.2561 | < 0.0001 |
| ti(Trial.s,Lat.XY):Typeflexible | 4.4415 | 16.0000 | 1.3359 | < 0.0001 |
| ti(Trial.s,Sag.XY):Typeflexible | 2.6761 | 16.0000 | 1.5412 | < 0.0001 |
| ti(Lat.XY,Sag.XY):Typeflexible | 0.0017 | 17.0000 | 0.0001 | 0.4377 |
| ti(Trial.s,Lat.XY,Sag.XY) | 0.0008 | 68.0000 | 0.0000 | 0.4034 |
| ti(Trial.s,Lat.XY,Sag.XY):Typeflexible | 0.0005 | 68.0000 | 0.0000 | 1.0000 |
| s(tpnt.s,Subject) | 288.5103 | 314.0000 | 566.1167 | < 0.0001 |
| s(Type,Subject) | 33.6190 | 67.0000 | 14.0516 | < 0.0001 |

### 3 A3. Theta Power (IRASA)

| A. parametric coefficients | Estimate | Std. Error | t-value | p-value |
|---|---|---|---|---|
| (Intercept) | 0.0928 | 0.0369 | 2.5171 | 0.0118 |
| bln | 0.1120 | 0.0030 | 37.7802 | $< 0.0001$ |
| Typeflexible | -0.0110 | 0.0141 | -0.7772 | 0.4371 |
| B. smooth terms | edf | Ref.df | F-value | p-value |
| ti(Trial.s) | 3.6568 | 4.0000 | 19.1232 | $< 0.0001$ |
| ti(Lat.XY) | 3.7736 | 4.0000 | 302.2238 | $< 0.0001$ |
| ti(Sag.XY) | 3.9309 | 4.0000 | 1844.7768 | $< 0.0001$ |
| ti(Trial.s):Typeflexible | 0.8236 | 4.0000 | 1.2524 | 0.0135 |
| ti(Lat.XY):Typeflexible | 0.8283 | 4.0000 | 1.1892 | 0.0032 |
| ti(Sag.XY):Typeflexible | 0.0007 | 4.0000 | 0.0001 | 0.5886 |
| ti(Trial.s,Lat.XY) | 1.5748 | 16.0000 | 0.4126 | 0.0035 |
| ti(Trial.s,Sag.XY) | 2.4140 | 15.0000 | 1.3467 | $< 0.0001$ |
| ti(Lat.XY,Sag.XY) | 16.6383 | 17.0000 | 32.9801 | $< 0.0001$ |
| ti(Trial.s,Lat.XY):Typeflexible | 3.5371 | 16.0000 | 0.5128 | 0.0098 |
| ti(Trial.s,Sag.XY):Typeflexible | 0.0018 | 16.0000 | 0.0001 | 0.3715 |
| ti(Lat.XY,Sag.XY):Typeflexible | 5.7300 | 17.0000 | 1.6929 | $< 0.0001$ |
| ti(Trial.s,Lat.XY,Sag.XY) | 0.0007 | 68.0000 | 0.0000 | 0.4258 |
| ti(Trial.s,Lat.XY,Sag.XY):Typeflexible | 7.6258 | 68.0000 | 0.2204 | 0.0026 |
| s(tpnt.s,Subject) | 291.4877 | 314.0000 | 1393.1282 | $< 0.0001$ |
| s(Type,Subject) | 33.2875 | 67.0000 | 16.4953 | $< 0.0001$ |

### 4 A4. Alpha Power (IRASA)

| A. parametric coefficients | Estimate | Std. Error | t-value | p-value |
|---|---|---|---|---|
| (Intercept) | 0.1956 | 0.0481 | 4.0634 | < 0.0001 |
| bln | 0.3083 | 0.0034 | 91.9256 | < 0.0001 |
| Typeflexible | -0.0116 | 0.0135 | -0.8588 | 0.3904 |
| B. smooth terms | edf | Ref.df | F-value | p-value |
| ti(Trial.s) | 3.2032 | 4.0000 | 12.4987 | 0.0001 |
| ti(Lat.XY) | 3.7409 | 4.0000 | 181.6516 | < 0.0001 |
| ti(Sag.XY) | 3.9574 | 4.0000 | 185.6785 | < 0.0001 |
| ti(Trial.s):Typeflexible | 3.2042 | 4.0000 | 13.4623 | < 0.0001 |
| ti(Lat.XY):Typeflexible | 0.1619 | 4.0000 | 0.0482 | 0.1683 |
| ti(Sag.XY):Typeflexible | 1.4308 | 4.0000 | 1.1028 | 0.0185 |
| ti(Trial.s,Lat.XY) | 2.1258 | 16.0000 | 0.2403 | 0.0552 |
| ti(Trial.s,Sag.XY) | 9.1852 | 16.0000 | 3.0641 | < 0.0001 |
| ti(Lat.XY,Sag.XY) | 15.9011 | 17.0000 | 75.1577 | < 0.0001 |
| ti(Trial.s,Lat.XY):Typeflexible | 1.6442 | 16.0000 | 0.2412 | 0.0495 |
| ti(Trial.s,Sag.XY):Typeflexible | 0.0012 | 16.0000 | 0.0000 | 0.7115 |
| ti(Lat.XY,Sag.XY):Typeflexible | 2.2677 | 17.0000 | 0.3494 | 0.0116 |
| ti(Trial.s,Lat.XY,Sag.XY) | 1.2087 | 68.0000 | 0.0260 | 0.1052 |
| ti(Trial.s,Lat.XY,Sag.XY):Typeflexible | 0.3672 | 68.0000 | 0.0060 | 0.2295 |
| s(tpnt.s,Subject) | 298.6820 | 314.0000 | 5592.8961 | < 0.0001 |
| s(Type,Subject) | 32.9176 | 67.0000 | 21.5883 | < 0.0001 |

### 5 A5. Beta Power (IRASA)

| A. parametric coefficients | Estimate | Std. Error | t-value | p-value |
|---|---|---|---|---|
| (Intercept) | -0.2349 | 0.0310 | -7.5895 | < 0.0001 |
| bln | 0.4704 | 0.0026 | 181.1611 | < 0.0001 |
| Typeflexible | 0.0187 | 0.0109 | 1.7143 | 0.0865 |
| B. smooth terms | edf | Ref.df | F-value | p-value |
| ti(Trial.s) | 2.8009 | 4.0000 | 15.2897 | < 0.0001 |
| ti(Lat.XY) | 2.9640 | 4.0000 | 68.5055 | < 0.0001 |
| ti(Sag.XY) | 2.8927 | 4.0000 | 133.4812 | < 0.0001 |
| ti(Trial.s):Typeflexible | 2.9369 | 4.0000 | 30.0801 | < 0.0001 |
| ti(Lat.XY):Typeflexible | 0.2356 | 4.0000 | 0.0714 | 0.2737 |
| ti(Sag.XY):Typeflexible | 0.8932 | 4.0000 | 2.0855 | 0.0021 |
| ti(Trial.s,Lat.XY) | 7.6330 | 16.0000 | 2.1561 | < 0.0001 |
| ti(Trial.s,Sag.XY) | 8.1742 | 16.0000 | 3.4591 | < 0.0001 |
| ti(Lat.XY,Sag.XY) | 16.5932 | 17.0000 | 33.0392 | < 0.0001 |
| ti(Trial.s,Lat.XY):Typeflexible | 6.8156 | 16.0000 | 1.2368 | 0.0002 |
| ti(Trial.s,Sag.XY):Typeflexible | 0.0006 | 16.0000 | 0.0000 | 0.5986 |
| ti(Lat.XY,Sag.XY):Typeflexible | 0.0015 | 17.0000 | 0.0001 | 0.4915 |
| ti(Trial.s,Lat.XY,Sag.XY) | 4.6218 | 68.0000 | 0.0981 | 0.0543 |
| ti(Trial.s,Lat.XY,Sag.XY):Typeflexible | 0.0025 | 68.0000 | 0.0000 | 0.3830 |
| s(tpnt.s,Subject) | 298.7249 | 314.0000 | 1117.9927 | < 0.0001 |
| s(Type,Subject) | 32.8490 | 67.0000 | 13.0557 | < 0.0001 |

Table 1: 1/f and delta power on learning (IRASA)

| | Chisq | Df | Pr(>Chisq) |
|---|---|---|---|
| delta_dif_log | 0.0638651 | 1 | 0.8004881 |
| dif_bexp | 0.9466432 | 1 | 0.3305752 |
| type | 23.7084776 | 1 | 0.0000011 |
| sag | 0.0785787 | 2 | 0.9614725 |
| lat | 0.0046301 | 2 | 0.9976876 |
| delta_dif_log:dif_bexp | 0.1294122 | 1 | 0.7190422 |
| delta_dif_log:type | 20.7774507 | 1 | 0.0000052 |
| dif_bexp:type | 27.4417905 | 1 | 0.0000002 |
| delta_dif_log:sag | 0.1016205 | 2 | 0.9504590 |
| dif_bexp:sag | 1.6040889 | 2 | 0.4484113 |
| type:sag | 0.1541287 | 2 | 0.9258303 |
| delta_dif_log:lat | 0.3642174 | 2 | 0.8335108 |
| dif_bexp:lat | 0.3429524 | 2 | 0.8424203 |
| type:lat | 3.9184956 | 2 | 0.1409644 |
| sag:lat | 0.1347929 | 4 | 0.9978284 |
| delta_dif_log:dif_bexp:type | 12.4761050 | 1 | 0.0004122 |
| delta_dif_log:dif_bexp:sag | 1.7976682 | 2 | 0.4070440 |
| delta_dif_log:type:sag | 0.8112448 | 2 | 0.6665618 |
| dif_bexp:type:sag | 0.2676567 | 2 | 0.8747402 |
| delta_dif_log:dif_bexp:lat | 1.2124644 | 2 | 0.5454020 |
| delta_dif_log:type:lat | 1.5923322 | 2 | 0.4510550 |
| dif_bexp:type:lat | 1.3333536 | 2 | 0.5134119 |
| delta_dif_log:sag:lat | 0.0917976 | 4 | 0.9989783 |
| dif_bexp:sag:lat | 0.1553944 | 4 | 0.9971335 |
| type:sag:lat | 0.3972499 | 4 | 0.9827014 |
| delta_dif_log:dif_bexp:type:sag | 2.5330786 | 2 | 0.2818052 |
| delta_dif_log:dif_bexp:type:lat | 2.9380684 | 2 | 0.2301477 |
| delta_dif_log:dif_bexp:sag:lat | 1.0106783 | 4 | 0.9081725 |
| delta_dif_log:type:sag:lat | 0.4538530 | 4 | 0.9778351 |
| dif_bexp:type:sag:lat | 1.3190546 | 4 | 0.8581344 |
| delta_dif_log:dif_bexp:type:sag:lat | 1.0251953 | 4 | 0.9059518 |

Table 1: 1/f and theta power on learning (IRASA)

| | Chisq | Df | Pr(>Chisq) |
|---|---|---|---|
| theta_dif_log | 0.0031288 | 1 | 0.9553933 |
| dif_bexp | 0.3737973 | 1 | 0.5409417 |
| type | 22.4196746 | 1 | 0.0000022 |
| sag | 0.0475055 | 2 | 0.9765271 |
| lat | 0.0066842 | 2 | 0.9966635 |
| theta_dif_log:dif_bexp | 0.0002710 | 1 | 0.9868648 |
| theta_dif_log:type | 4.5774379 | 1 | 0.0323956 |
| dif_bexp:type | 17.4299930 | 1 | 0.0000298 |
| theta_dif_log:sag | 0.0651907 | 2 | 0.9679302 |
| dif_bexp:sag | 1.0460229 | 2 | 0.5927329 |
| type:sag | 0.0989554 | 2 | 0.9517264 |
| theta_dif_log:lat | 0.0221312 | 2 | 0.9889954 |
| dif_bexp:lat | 0.0394554 | 2 | 0.9804656 |
| type:lat | 1.6183010 | 2 | 0.4452361 |
| sag:lat | 0.0139291 | 4 | 0.9999759 |
| theta_dif_log:dif_bexp:type | 2.3900954 | 1 | 0.1221062 |
| theta_dif_log:dif_bexp:sag | 0.0403813 | 2 | 0.9800118 |
| theta_dif_log:type:sag | 0.3964568 | 2 | 0.8201825 |
| dif_bexp:type:sag | 0.0095446 | 2 | 0.9952391 |
| theta_dif_log:dif_bexp:lat | 0.0004885 | 2 | 0.9997558 |
| theta_dif_log:type:lat | 0.1294694 | 2 | 0.9373161 |
| dif_bexp:type:lat | 0.9937995 | 2 | 0.6084140 |
| theta_dif_log:sag:lat | 0.3506557 | 4 | 0.9863138 |
| dif_bexp:sag:lat | 0.0282033 | 4 | 0.9999015 |
| type:sag:lat | 0.0360091 | 4 | 0.9998399 |
| theta_dif_log:dif_bexp:type:sag | 1.7957661 | 2 | 0.4074313 |
| theta_dif_log:dif_bexp:type:lat | 0.8246814 | 2 | 0.6620986 |
| theta_dif_log:dif_bexp:sag:lat | 0.9341132 | 4 | 0.9196165 |
| theta_dif_log:type:sag:lat | 0.1867883 | 4 | 0.9959010 |
| dif_bexp:type:sag:lat | 0.8318502 | 4 | 0.9341278 |
| theta_dif_log:dif_bexp:type:sag:lat | 0.5244378 | 4 | 0.9710788 |

Table 1: 1/f and alpha power on learning (IRASA)

| | Chisq | Df | Pr(>Chisq) |
|---|---|---|---|
| alpha_dif_log | 0.0001376 | 1 | 0.9906418 |
| dif_bexp | 0.7075997 | 1 | 0.4002418 |
| type | 24.7733601 | 1 | 0.0000006 |
| sag | 0.0050041 | 2 | 0.9975011 |
| lat | 0.0417828 | 2 | 0.9793253 |
| alpha_dif_log:dif_bexp | 6.7456337 | 1 | 0.0093977 |
| alpha_dif_log:type | 1.0999706 | 1 | 0.2942726 |
| dif_bexp:type | 5.8647679 | 1 | 0.0154469 |
| alpha_dif_log:sag | 1.3907574 | 2 | 0.4988855 |
| dif_bexp:sag | 0.3660057 | 2 | 0.8327658 |
| type:sag | 0.5591930 | 2 | 0.7560888 |
| alpha_dif_log:lat | 0.1468749 | 2 | 0.9291943 |
| dif_bexp:lat | 0.4503652 | 2 | 0.7983704 |
| type:lat | 0.3247628 | 2 | 0.8501169 |
| sag:lat | 0.4397158 | 4 | 0.9790980 |
| alpha_dif_log:dif_bexp:type | 15.9426120 | 1 | 0.0000653 |
| alpha_dif_log:dif_bexp:sag | 3.6813224 | 2 | 0.1587125 |
| alpha_dif_log:type:sag | 0.2552763 | 2 | 0.8801718 |
| dif_bexp:type:sag | 1.4777658 | 2 | 0.4776472 |
| alpha_dif_log:dif_bexp:lat | 0.8548861 | 2 | 0.6521745 |
| alpha_dif_log:type:lat | 0.1879070 | 2 | 0.9103251 |
| dif_bexp:type:lat | 2.2262512 | 2 | 0.3285305 |
| alpha_dif_log:sag:lat | 0.3684978 | 4 | 0.9849739 |
| dif_bexp:sag:lat | 1.3776789 | 4 | 0.8480651 |
| type:sag:lat | 1.2392577 | 4 | 0.8715938 |
| alpha_dif_log:dif_bexp:type:sag | 0.2231206 | 2 | 0.8944375 |
| alpha_dif_log:dif_bexp:type:lat | 3.4887781 | 2 | 0.1747517 |
| alpha_dif_log:dif_bexp:sag:lat | 0.2338203 | 4 | 0.9936760 |
| alpha_dif_log:type:sag:lat | 0.7199022 | 4 | 0.9488521 |
| dif_bexp:type:sag:lat | 1.4243033 | 4 | 0.8399602 |
| alpha_dif_log:dif_bexp:type:sag:lat | 3.1143612 | 4 | 0.5388730 |

Table 1: 1/f and beta power on learning (IRASA)

| | Chisq | Df | Pr(>Chisq) |
|---|---|---|---|
| beta_dif_log | 3.3786624 | 1 | 0.0660456 |
| dif_bexp | 1.0055650 | 1 | 0.3159677 |
| type | 25.7741827 | 1 | 0.0000004 |
| sag | 0.0582083 | 2 | 0.9713153 |
| lat | 0.0718273 | 2 | 0.9647236 |
| beta_dif_log:dif_bexp | 0.0910310 | 1 | 0.7628705 |
| beta_dif_log:type | 45.8228655 | 1 | 0.0000000 |
| dif_bexp:type | 22.2866853 | 1 | 0.0000023 |
| beta_dif_log:sag | 0.1487802 | 2 | 0.9283095 |
| dif_bexp:sag | 0.6215698 | 2 | 0.7328715 |
| type:sag | 0.0652476 | 2 | 0.9679026 |
| beta_dif_log:lat | 0.1369093 | 2 | 0.9338358 |
| dif_bexp:lat | 0.2467065 | 2 | 0.8839513 |
| type:lat | 0.2109777 | 2 | 0.8998845 |
| sag:lat | 0.3857642 | 4 | 0.9836257 |
| beta_dif_log:dif_bexp:type | 30.9696876 | 1 | 0.0000000 |
| beta_dif_log:dif_bexp:sag | 0.0237201 | 2 | 0.9882100 |
| beta_dif_log:type:sag | 2.3243458 | 2 | 0.3128058 |
| dif_bexp:type:sag | 0.4975626 | 2 | 0.7797505 |
| beta_dif_log:dif_bexp:lat | 0.4672160 | 2 | 0.7916721 |
| beta_dif_log:type:lat | 0.3725366 | 2 | 0.8300508 |
| dif_bexp:type:lat | 2.2410968 | 2 | 0.3261009 |
| beta_dif_log:sag:lat | 0.9887833 | 4 | 0.9114920 |
| dif_bexp:sag:lat | 0.1700010 | 4 | 0.9965858 |
| type:sag:lat | 1.6213721 | 4 | 0.8049459 |
| beta_dif_log:dif_bexp:type:sag | 1.5213623 | 2 | 0.4673480 |
| beta_dif_log:dif_bexp:type:lat | 0.8599960 | 2 | 0.6505104 |
| beta_dif_log:dif_bexp:sag:lat | 0.3628628 | 4 | 0.9854029 |
| beta_dif_log:type:sag:lat | 0.9887146 | 4 | 0.9115024 |
| dif_bexp:type:sag:lat | 1.1326172 | 4 | 0.8890632 |
| beta_dif_log:dif_bexp:type:sag:lat | 0.5050094 | 4 | 0.9730115 |
